# Augmenting complex and dynamic performance through mindfulness-based cognitive training: an investigation of on-task EEG dynamics

**DOI:** 10.1101/2024.07.29.605541

**Authors:** Chloe A. Dziego, Ina Bornkessel-Schlesewsky, Maarten A. Immink, Zachariah R. Cross, Matthias Schlesewsky

**Affiliations:** Cognitive Neuroscience Laboratory, Australian Research Centre for Interactive and Virtual Environments, University of South Australia, Adelaide, Australia; Sport, Health, Activity, Performance and Exercise (SHAPE) Research Centre, Flinders University, Adelaide, Australia; Dynamic Brain Lab, Department of Medical Social Sciences, Northwestern Feinberg School of Medicine, Chicago, Illinois, United States of America

**Keywords:** mindfulness, cognitive training, EEG, alpha, theta, aperiodic

## Abstract

Mindfulness-based cognitive training exhibits great propensity for improving cognitive performance across a range of contexts. However, the neurophysiological basis of these cognitive enhancements has remained relatively unclear. Previous studies have widely examined EEG *during* mindfulness practice – or made comparisons with long-term meditators and controls – but have failed to capture how EEG dynamics in *subsequent* cognitive testing scenarios might be altered as a function of mindfulness-based interventions. The current study therefore aimed to assess a variety of EEG dynamics (oscillatory, aperiodic, and event-related) during engagement in a dynamic and complex cognitive task, following a mindfulness-based cognitive training regime. Participants (n = 40, age range = 18 – 38) attended the lab on two separate occasions (pre- and post-a web-based one-week mindfulness intervention), where EEG was recorded during engagement in the Target Motion Analyst (TMA) task. Previous analysis of the same participants demonstrated that greater adherence to the mindfulness-based cognitive training was associated with improved performance on the TMA task (Dziego, Bornkessel-Schlesewsky, Schlesewsky et al., 2024). Here, we capitalise on these previous findings to assess whether adherence is paralleled by measurable differences in on-task EEG dynamics. Linear mixed-effects modelling demonstrated that, while main effects were observed across session, adherence to cognitive training was not directly associated with alpha power, theta power or 1/*f* parameters. Challenges also arose when computing event-related potentials (ERPs), illustrating the difficulties of using this technique in more complex testing environments. While these results are challenging to place within the context of previous EEG studies on meditation and cognitive performance, our findings highlight the complexities in understanding the cognitive benefits of mindfulness-based training interventions through EEG dynamics observed during subsequent cognitive testing.

## 1. Introduction

While mindfulness techniques are commonly used to enhance psychological health and wellbeing (Keng et al., 2011; Khoury et al., 2013), human performance applications are demonstrating how mindfulness practices can further facilitate improved cognition and performance, producing transferable enhancements across a range of contexts (Ainsworth et al., 2013; Brown et al., 2022; Dziego, Bornkessel-Schlesewsky, Schlesewsky, et al., 2024; Jha et al., 2015). While theories have been developed about the underlying mechanisms of cognitive enhancements from mindfulness (i.e., enhancing attentional capacity; Gallant, 2016; Malinowski, 2013, or improving memory consolidation; Brown et al., 2022; Dziego, Bornkessel-Schlesewsky, Schlesewsky et al., 2024) and many studies have observed diverse neural correlates (Cahn & Polich, 2006; Edwards et al., 2012; Lomas et al., 2015; Tang et al., 2015), there is little consensus on the exact neurophysiological mechanisms underlying the observed performance outcomes. Explorations have largely focused on assessing neural correlates *during* mindfulness engagement or comparing long-term practitioners with controls (as described in Cahn & Polich, 2006), limiting our knowledge of how EEG dynamics during information processing may be modified as a function of mindfulness training. Very few studies have explicitly examined cognitive benefits through the perspective of on-task neurophysiological change (i.e., changes in the brain’s activity *during* cognitive testing), which may help to explain performance gains over and above other more common analysis approaches (e.g., in the resting-state or during mindfulness practice). With this in mind, the current study aims to examine on-task EEG activity (i.e., through oscillatory, aperiodic and event-related potential methods) following engagement in a one-week mindfulness-based cognitive training regime. Herein, we aim to elucidate how brain changes from mindfulness training – associated with improved cognitive functioning – are exhibited in on-task EEG activity.

### 1.1 Oscillatory Correlates of Meditation

As described above, the extant literature surrounding EEG correlates of mindfulness and meditation has largely focused on measuring individuals *during* mindfulness engagement or comparing long-term practitioners with controls. Current perspectives broadly suggest that engagement in meditation is most often associated with increases in power and synchronisation of both alpha and theta frequencies within EEG signals (see reviews, Cahn & Polich, 2006; Lee et al., 2018; Lomas et al., 2015). In particular, alpha power is reported to increase during meditation, when compared to control conditions, for both new and experienced meditators (Ahani et al., 2014; Bing-Canar et al., 2016; Huang & Lo, 2009; Lagopoulos et al., 2009; Takahashi et al., 2005; Tsai et al., 2013). Topographically, these increases have been observed in frontal (Takahashi et al., 2005) and posterior regions (Ahani et al., 2014; Cahn et al., 2010; Dentico et al., 2018; Lagopoulos et al., 2009; Lee et al., 2018) and parallel findings within fMRI-PET studies (whereby increased alpha power is related to decreased cortical blood flow; Goldman et al., 2002). On the other hand, increased theta power is often noted, again, with individuals both new to and experienced with meditation (Ahani et al., 2014; Dentico et al., 2018;

Lomas et al., 2014; Takahashi et al., 2005) during present engagement, when compared to signals recorded at-rest. Similarly, this increase has been mostly documented in frontal regions (Dentico et al., 2018; Lee et al., 2018; Lomas et al., 2014; Takahashi et al., 2005; Tanaka et al., 2014), although other studies have also reported increases across the cortex (e.g., Ahani et al., 2014). However, one review notes that increases in theta are more often exhibited by experienced, when compared to novel, practitioners (Lomas et al., 2015). Additionally, as relationships are also exhibited between alpha, theta and non-meditative tasks (e.g., in mind-wandering; Compton et al., 2010 or task engagement; Benedek et al., 2014; Klimesch, 1999), it is difficult to confidently link these changes exclusively to mindfulness engagement. Likewise, fluctuations in alpha power can be related to a range of relaxation techniques (Morse et al., 1977; Teplan et al., 2014) and other authors have suggested that these correlates are related to acute changes from concentration; not necessarily directly stemming from engagement in meditation (Tanaka et al., 2014).

On the other hand, when exploring EEG correlates more longitudinally, results similarly point to key differences in alpha and theta power during the resting-state when experienced meditators are compared with novel practitioners or non-meditating controls (Cahn & Polich, 2006). Here, it is believed that multiple inductions of the mindful state evoke more persistent change to individuals’ trait levels (Cahn & Polich, 2006; Kiken et al., 2015). Consistent with this, studies have observed increased alpha (Aftanas & Golosheykin, 2005; Corby et al., 1978; Kasamatsu & Hirai, 1966; Travis et al., 2002) and theta (Aftanas & Golosheykin, 2005; Corby et al., 1978; Tanaka et al., 2014) amplitudes during the resting-state for long-term meditators versus control participants. However, some authors report that EEG is not altered further with longer-term engagement in mindfulness practices (Lomas et al., 2014; Lutz et al., 2004), highlighting how more trait-like cognitive changes could instead be a direct function of current practice (Lodha & Gupta, 2022; Teper & Inzlicht, 2013).

Nevertheless, Fell and colleagues (2010) consider alpha and theta synchronisation the key ‘signatures’ of meditation practices due to their ubiquitous nature across styles, general experience and practice depth. There is also clear alignment between the functional correlates of alpha and theta (explored below) and the subjective experience of mindfulness/meditation more generally; the co-presence of these oscillatory signals during mindfulness are thought to relate to a state of ‘relaxed-alertness’ (Britton et al., 2014). While there have been few contrasting findings in the area (e.g. no difference or decreases in alpha; Amihai & Kozhevnikov, 2014; Cahn et al., 2010; Tsai et al., 2013 or theta power; Huang & Lo, 2009; Milz et al., 2014) and some relationships have been observed across alternative oscillatory frequencies (e.g., gamma; Cahn et al., 2010; Dentico et al., 2018; Lutz et al., 2004 or beta; Ahani et al., 2014), there is overall a large consensus within the literature that alpha and theta power are clear correlates of meditation engagement. However, previous EEG findings have rarely been considered in tandem with the salutary benefits of mindfulness or contextualised within the larger literature of cognitive performance (c.f., Dziego, Bornkessel-Schlesewsky, Schlesewsky et al., 2024). Thus, there is little understanding of *how* mindfulness training impacts neurophysiological states captured on-task (external to direct engagement or in the resting-state) and consequently, how this might relate to enhanced information processing capabilities.

### 1.2 Oscillatory and Aperiodic Correlates of Cognitive Performance

Research focusing on individual cognitive performance exhibits results that align well with previous findings surrounding the EEG correlates of meditation. For example, while increased alpha activity was once considered a correlate of the brain’s ‘idling state’, it is now viewed as a key component of higher-order cognition (Bazanova & Vernon, 2014). It is believed that alpha oscillations are associated with the inhibition of task-irrelevant cortical areas, which in turn may facilitate improved performance in cognitive testing (Klimesch et al., 2007). Consistently, theoretical explanations describe alpha as a mechanism for increasing signal to noise ratio by inhibiting conflicting processes (Cooper et al., 2003; von Stein & Sarnthein, 2000). In line with this, studies have shown that increases in alpha activity during task engagement relate to improved performance on semantic long-term memory tasks (Klimesch, 1999) and mental rotation tasks (Cooper et al., 2003) and can reflect the strength of attentional biasing (Kelly et al., 2009; Thut et al., 2006). On the other hand, theta oscillations have been broadly linked to multiple cognitive processes including attentional switching (Dietl et al., 1999) and spatial navigation (Korotkova et al., 2018) but are critically noted to play a role in memory formation and retrieval (e.g., Berry & Thompson, 1978; Klimesch, 1999; Klimesch et al., 1997; Seager et al., 2002). Here, increases in theta power (i.e., greater synchronisation) have been consistently related to cognitive performance in a variety of memory paradigms (e.g., Griffin et al., 2004; Herweg et al., 2020; Klimesch et al., 1997; Seager et al., 2002). Alternative perspectives detail that it is not just increases in alpha or theta power alone that predict cognitive performance, but a dynamic interplay between the two frequencies that facilitate improved information processing (e.g., Klimesch, 1999; Raufi & Longo, 2022).

Despite the promising implications of this activity, studies aiming to directly uncover the underlying mechanisms behind improvements to cognition from mindfulness-based cognitive training have rarely considered these two components (i.e., the neural correlates of mindfulness and separably, cognitive performance) in parallel. In investigations of mindfulness-induced performance-related EEG change, only a handful of studies have examined oscillatory measures directly within subsequent cognitive testing scenarios. For example, after a short mindful breathing induction, Bing-Canar et al. (2016) demonstrated that participants exhibited greater error-related alpha suppression during a Stroop task. These findings are interpreted to represent enhanced self- and error-monitoring, in addition to a more dynamic control of attention during task engagement. Similarly, Wong et al. (2018) observed that alpha power desynchronisation (during a psychomotor vigilance task) was reduced with greater attendance to an 8-week mindfulness training intervention. Here, higher adherence to the mindfulness training was related to improved ability to suppress background cortical activity (Wong et al., 2018).

Altogether, these findings suggest that both alpha and theta activity may prove useful in capturing the cognitive benefits of mindfulness training.

In addition to this, studies are now demonstrating that scale-free neural activity, known as aperiodic 1/*f* activity, may play a critical role in cognitive function (Donoghue et al., 2020). The examination of aperiodic 1/*f* activity is becoming more prominent within the EEG literature, with investigations exploring its relationship to states of consciousness (e.g., Chatburn et al., 2024; Colombo et al., 2019; Lendner et al., 2020), cognition (Cross, Corcoran, et al., 2022; Dave et al., 2018; Dziego et al., 2023; Immink et al., 2021; Ouyang et al., 2020; Sheehan et al., 2018) and psychiatric disorders (Ostlund et al., 2021; Peterson et al., 2017). Aperiodic activity manifests as a quasi-straight line through power spectral density (PSD) plots, demonstrating the 1/*f* power law (whereby lower frequencies typically exhibit higher power, and vice versa; He, 2014; Kayser & Ermentrout, 2010). While some resting-state meditation studies have begun to consider this EEG component (Dziego, Bornkessel-Schlesewsky, Schlesewsky et al., 2024; Rodriguez-Larios et al., 2021), many previous oscillatory investigations during meditation have completely ignored underlying 1/*f* activity. Critically, we hypothesise that changes from meditation (explored above) may be better explained through an ‘exponent change’ (i.e., where the regression line through the PSD tilts at a centre axis; Donoghue et al., 2020), as opposed to changes in theta/alpha. Donoghue et al. (2020, pg. 1656) remarks that “failing to consider aperiodic activity confounds oscillatory measures, and masks crucial behaviourally and physiologically relevant information”. In this case, we aim to also consider aperiodic 1/*f* activity within the current study through multiple lenses: (1) as a behaviourally relevant phenomenon (related to neurophysiological change with engagement in mindfulness) and (2) as a confounding variable to be considered when estimating oscillatory measures.

### 1.3 Event-Related Potential Investigations

In addition to oscillatory or time-frequency analysis, another popular approach within EEG investigations of cognitive performance are event-related potentials (ERPs). Event-related potentials provide temporally fine-grained insight into stimulus-locked activity averaged across experimental trials (Luck, 2014), which aim to capture microvolt (mV) change prior to and succeeding external stimulation, task execution or other cognitive events. As such, ERPs have been employed in a range of inquiries into long-term neuroplastic change from mindfulness practice and, in comparison to oscillatory studies, have often been better positioned to capture how cognitive benefits arise (due to their evocation *during* cognitive testing or paradigm engagement).

In the mindfulness literature, comparisons of experienced meditators and controls demonstrate key differences in brainstem (e.g., McEvoy et al., 1980), middle latency (e.g., Telles et al., 1993, 1994; Telles & Naveen, 2004) and early sensory potentials (e.g., Kobal et al., 1975; Zhang et al., 1993) which can theoretically be linked to improvements in sensory and attentional processes (for a review, see Cahn & Polich, 2006). In studies specifically investigating the cognitive effects of meditation, the P300 and Contingent Negative Variation (CNV) have garnered significant interest. Studies have shown that meditative focusing is associated with increased P300 amplitudes and reduced latency in a traditional auditory oddball task, which the authors related to enhanced attentional processes (Delgado-Pastor et al., 2013; Telles et al., 2019). Wong et al. (2018) also examined changes in the P300 in parallel with performance on a psychomotor vigilance task following an 8-week mindfulness training intervention. Here, performance decrements seen over a high-demand period were mitigated by attendance to the mindfulness training program but, while P300 amplitude reductions were less pronounced across time-on-task (when compared to controls), these effects were not significant. Yoshida et al. (2020) examined the P300 during an oddball task following an 8-week focused attention meditation intervention. Their findings similarly demonstrated that P300 amplitudes were higher to oddball stimuli following mindfulness training, paralleling faster response times. Other studies have also observed reduced P300 amplitudes to task-irrelevant stimuli (when compared to controls), associated with improvements to top-down inhibition (Cahn & Polich, 2009; Moore et al., 2012). On the other hand, more pronounced amplitudes of the CNV (related to anticipatory processes) following mindfulness-based interventions have been observed across multiple studies (e.g., Incagli et al., 2020; Paty et al., 1978; Travis et al., 2002) in both state-induced and more trait-like fashions (Cahn & Polich, 2006). In comparison to oscillatory measures (occurring across larger timescales), previous work demonstrates that ERPs provide a promising avenue for assessing more minute information processing differences and could be useful in assessing neurophysiological change associated with enhanced cognition following mindfulness training.

### 1.4 The Current Study

Overall, the extant literature on meditation has largely focused on EEG recordings *during* mindfulness, which do not provide insight into more persisting change that might help to explain cognitive enhancement. Here, we highlight a critical gap within prior investigations: there is a lack of understanding regarding the traceable changes garnered from mindfulness training that relate to functional changes underlying cognitive enhancements (observed outside of the practice itself). In the oscillatory domain, there are only a handful of studies assessing neurophysiological change in tandem with performance enhancements following mindfulness (e.g., Bing-Canar et al., 2016 and Wong et al., 2018). While these studies suggest that alpha power may play a role in subsequent cognitive enhancements, there is still great room to expand on investigations in the oscillatory domain. Moreover, previous ERP investigations have captured how sensory and attentional processing may be enhanced by mindfulness engagement, however, these studies have been limited to traditional cognitive tests (e.g., Delgado-Pastor et al., 2013; Wong et al., 2018; Yoshida et al., 2020). The current study aims to characterise on-task EEG signatures of mindfulness-based cognitive training, capitalising on previously observed performance enhancements in a more complex and dynamic cognitive task (Dziego, Bornkessel-Schlesewsky, Schlesewsky et al., 2024). Our study is particularly innovative in exploring performance in a novel testing scenario and assessing mindfulness adherence in an individualised dose-dependent fashion (versus commonplace ‘intervention versus control group’ designs; see Dziego, Bornkessel-Schlesewsky, Schlesewsky et al., 2024). In this way, the current study aims to assess how brain changes from mindfulness may be a direct function of current practice (as has been suggested of the cognitive benefits; Lodha & Gupta, 2022; Teper & Inzlicht, 2013) and exact repetitions of induced mindful states (Cahn & Polich, 2006; Kiken et al., 2015).

We aim to approach this question through the exploration of a range of oscillatory (alpha and theta power), aperiodic (1/*f*) and event-related potential EEG dynamics during cognitive testing. We hypothesise that higher adherence to the mindfulness-based cognitive training regime will be related to increased alpha and theta power during subsequent testing, aligning with both the mindfulness and cognitive performance literature more broadly. Moreover, while engaging in mindfulness practice is known to influence inhibitory neurotransmitters within the brain (Edwards et al., 2012; Elias et al., 2000) and has been related to steeper 1/*f* slopes (Rodriguez-Larios et al., 2021), we propose that flatter 1/*f* slopes during testing will be associated with adherence to the cognitive training regime (i.e., better performance). Here, our hypothesis aligns with previous cognitive performance literature demonstrating that flatter 1/*f* slopes are superior in more dynamic contexts (Dziego et al., 2023) and where flatter slopes parallel superior performance after an 8-week mindfulness retreat (see Dziego, Zanesco, et al., 2024). Lastly, we conducted an exploratory analysis of event-related potentials around critical time-locked points of the cognitive testing session (e.g., when receiving auditory information) to assess differences in early sensory and subsequent components across levels of adherence.

## 2. Method

### 2.1 Participants

The current study included an analysis of EEG data recorded during Dziego, Bornkessel-Schlesewsky, Schlesewsky et al. (2024). The original study undertook a detailed analysis of performance augmentations and fluctuations in resting-state EEG measures, while the current study focuses on EEG data recorded during cognitive task engagement. The sample comprised of forty-two adults (M_age_ = 23.43, range: 18 – 38, females = 28) who volunteered to participate. Eligible participants reported no history of psychiatric, language, neurological or cognitive disorders. Consistent with conventional EEG studies, all participants were right-handed and did not report taking any medication that may affect their electrophysiology. Participants were eligible for the study only if they had no formal experience with alternative cognitive training or mindfulness programs. Participants received an honorarium of AUD$120 for participation in the study. Prior to the commencement of data collection, ethics approval was acquired from the University of South Australia’s ethics committee (project no. 204054). During data collection, two participants were able to attend only their first in-lab testing session (due to illness or other commitments) and were subsequently removed from the analysis. As a result, data from forty eligible participants were used in the final analysis (M_age_ = 23.50, range: 18 – 38, females = 26).

### 2.2 Mindfulness-Based Cognitive Training Intervention

The mindfulness-based cognitive training program consisted of seven distinct audio tracks (approximately 5 minutes in length) voiced by an experienced mindfulness instructor (Dr. M. Immink). Exercises guided participants through a range of focused attention and open monitoring mindfulness practices (for a review of types of mindfulness practice, see Crane et al., 2017; Lutz et al., 2008), including instructions to focus on breathing, being still, mentally scanning the body or externalising attention. All transcripts and audio recordings can be accessed via the Open Science Framework (OSF) repository here: https://osf.io/y9hvt/. The cognitive training program was accessible to participants through a webpage interface hosted on an online repository: Github Pages (GitHub Inc., 2021; accessible at https://github.com/chloeadee/cognitivetraining). *RMarkdown* in R Studio (R Core Team, 2020) was used to create the HTML code to produce the webpage interface. Participants were encouraged to listen and engage with the mindfulness exercises three times per day (for a total of 15 minutes per day). At the conclusion of their final session each day, participants were instructed to complete a ‘post-training’ questionnaire about their experience and report the number of sessions they were able to complete that day. These questions aimed to capture which day of training they had fulfilled (1-7), how many sessions were completed (0-3), and recorded how participants felt during their exercises. This data was, however, not used in the current analysis. Instead, objective completion rates were tracked using *Google Analytics* and *Google Tag Manager* (Google Inc., 2021). For a full description of the mindfulness-based cognitive training intervention and relevant questionnaires, see Dziego, Bornkessel-Schlesewsky, Schlesewsky et al. (2024).

### 2.3 Control-Room-Use-Simulation Environment

To capture performance in a more naturalistic and dynamic setting, we utilised the dual-screen Target Motion Analysis (TMA) task from within the Control-Room-Use-Simulation Environment (CRUSE; Michailovs et al., 2021). CRUSE is a medium-fidelity simulation incorporating multiple operator stations to mimic the roles within a submarine control room. While the simulation involves multiple potential operator stations, the current study examined individual performance, and thus focused only on the TMA task. Within a submarine, the TMA operator is responsible for creating a ‘tactical picture’ of the surrounding vessels (known as contacts) by integrating multiple sources of sensory information (see Dziego et al., 2023 and Dziego, Bornkessel-Schlesewsky, Schlesewsky et al., 2024 for a schematic of a participant engaging in the TMA task). Here, participants must integrate multiple sources of information from SONAR, optronics and the Track Manager to develop a ‘solution’ (i.e., an estimate of the location and movement) of each surrounding contact. For a more detailed description of the task, see Dziego et al. (2023). Performance within the task is measured by the distance between the coordinates of the participants’ plotted solutions and the simulation’s ‘truth.’ This metric is referred to as tactical picture error or TPE, whereby a lower score signifies better performance (i.e., less error). TPE is weighted by the priority of a contact (e.g., fishing vessel or warship), its course, range, and behaviour. For a detailed description of TPE, see Michailovs et al. (2021). At the commencement of the simulation, default solutions are included in the first recorded instances of TPE, with all participants starting at a score of 17,500. Throughout the simulation, TPE is logged in twenty second epochs. The TMA task also includes triggers around critical events within the simulation; that is, receiving information from SONAR, optics, and feedback from the track manager, as well as each time a participant ‘sets a solution.’ Within the current study, three different scenarios were used: (1) a practice scenario (involving two contacts with simple movement), (2) a pre- and (3) a post-training scenario. For comparable difficulty, both pre- and post-intervention simulations included five contacts (with more erratic movement) and were reflections and inversions of each other. The current analysis utilises TPE scores captured in sessions (2) and (3).

### 2.4 Protocol

Potential volunteers contacted researchers through email. To ensure eligibility, participants initially completed an online screening process before being provided access to an educational video and the TMA task. After accessing the video, participants were briefly questioned on their knowledge of the simulation to ensure they understood the critical components and general requirements of the task. Following this, participants booked two in-lab appointments (at the Cognitive Neuroscience Laboratory at the University of South Australia’s Magill campus) for their pre- and post-intervention testing sessions (same time, exactly one-week apart). Upon visiting the lab, participants provided written consent to participate before undergoing the EEG set up. Participants completed two minutes of resting-state EEG recording with eyes closed, before engaging in a practice scenario of the TMA task. The practice scenario (1) was 30 minutes in length whereby participants were instructed on the use of the simulation by the researcher (to see the script used for the practice session, see supplementary materials in Dziego et al., 2023). Participants were also able to clarify instructions and ask questions during this session. Following the practice scenario, participants engaged in a second TMA scenario (2), which ran for approximately 40 minutes. Here, participants were unable to receive feedback on their performance. EEG was continuously recorded throughout the 40-minute TMA scenario.

After their initial visit to the lab, participants were provided a unique link (via email) to access the mindfulness-based cognitive training intervention. Researchers encouraged participants to engage in the exercises three times per day for seven days (for a total of 15 minutes per day). After one week, participants attended their post-intervention testing session at the Cognitive Neuroscience Laboratory. The second testing session followed a comparable protocol to the first. Again, participants were fitted with an EEG cap, completed a two-minute resting-state recording (eyes closed) and engaged in a 40-minute testing session of the TMA task (3). In their second testing session, participants were not offered any further practice or feedback on their performance. EEG was recorded continuously throughout the 40-minute TMA scenario. The order of presentation of the two testing scenarios was randomised across participants.

## 3. Data Analysis

### 3.1 EEG recording and Pre-Processing

Continuous EEG was recorded throughout the TMA task from 32 Ag/AgCl electrodes mounted in an elastic cap (actiCAP, Brain Products GmbH, Gilching, Germany) through *OpenVibe* (Renard et al., 2010). To monitor eyeblinks, two electrodes were placed below and above the left eye. Channels were amplified using a LiveAmp 32 amplifier (Brain Products, GmbH) and sampled at a rate of 250 Hz. Impedances were kept below 10kΩ. All pre-processing and analysis of EEG data was performed using MNE-python (Gramfort et al., 2013). Due to unreliable scaling within the *OpenVibe* recording program, some participants’ data were rescaled (using the same scaling factor as conventional *BrainVision* recordings) to ensure comparable power estimates. Recordings requiring re-scaling were identified through visualisation of the raw EEG data PSD plots. Data were first re-referenced to TP9 and TP10 and PSD plots were visually screened to mark bad channels. EOG artifacts were identified and corrected using an Independent Component Analysis (ICA). Here, a copy of the raw data was bandpass filtered from 0.1 to 40 Hz. Independent Components were identified using the FastICA method (using EEG channels only) and epochs with a peak-to-peak voltages exceeding 250 microvolts were excluded. In MNE, the *create_eog_epochs()* and *ica.find_bads_eog()* functions were used to isolate EOG events and identify independent EOG components via correlation. Any identified components were subsequently removed from the original raw data. Bad channels were interpolated using the *interpolate_bads()* function. EEG data were then band-pass-filtered from 0.1 to 40 Hz (zero-phase, hamming windowed finite impulse response).

### 3.2 Aperiodic 1/*f* and Oscillatory Band Power Estimation

To estimate broadband and narrowband EEG changes across time, pre-processed EEG data were passed through the Irregular Resampling Auto-Spectral Analysis (IRASA) functions in the YASA toolbox (Vallat & Walker, 2021) in Python. This technique aims to separate oscillatory (periodic) signals from aperiodic (1/*f*) activity by iteratively resampling the power spectrum. This procedure allows the narrowband components to redistribute away from their original location, while the distribution of aperiodic activity remains constant. For a more detailed description of the IRASA method, see Wen & Liu (2016).

Pre-processed EEG data recorded during the TMA task were divided into 20 second epochs and passed through the *yasa.irasa()* function for estimation of broadband (aperiodic) and narrowband (oscillatory) parameters. We extracted individual aperiodic slope and intercept values across all electrodes for each 20 second epoch. In order to quantify oscillatory activity independent of the underlying aperiodic activity, mean power density across frequency bands was calculated from power estimates extracted from the *yasa.irasa()* function. These power estimates consider aperiodic activity by subtracting the mean regression fit of the aperiodic slope from the overall PSD. Within IRASA, power spectrums are calculated using the fast Fourier transform tapered with a Hanning window. This procedure was completed for all electrodes.

As recommended by Klimesch (2012), mean power densities were calculated according to individually tailored frequency ranges of each participant, in line with the harmonic frequency framework. This method suggests that frequency bands should be determined by each participants’ centre frequency (i.e., peak frequency within the alpha band), with upper and lower band bounds calculated using below formulae:

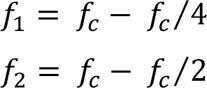

Here, *f*_c_ represents the centre frequency (also known as individual alpha frequency; IAF) and *f*_1_ and *f*_2_ indicate the lower and upper bands, respectively.

IAF estimates (Corcoran et al. 2018) were derived from 2 minutes of eyes-closed resting-state EEG recording prior to task engagement. Two-minute recordings were pre-processed using the same pipeline detailed above (excluding ICA fitting). The *philistine* package (Alday, 2019) in MNE-Python was used to calculate IAF estimates from parieto-occipital electrodes (P3/P4/O1/O2/Oz/P7/P9/Pz). This technique utilises a Savitsky-Golay filter (frame length = 11 frequency bins, polynomial degree = 5) to smooth the PSD before identifying peak activity within the alpha frequency (between 7 -13Hz). With each participant’s bounds for alpha and theta identified, power metrics for each band were calculated by averaging power estimates from each electrode across each individual frequency (e.g., 4 – 7Hz for theta) within the individually defined band range. For participants with bounds between frequencies (e.g., 4.2 Hz or 7.5Hz), bounds were rounded to the nearest whole number.

### 3.3 Event-Related Potentials

To examine event-related potentials around critical points within the TMA task, we epoched the processed EEG data around triggers of interest (i.e., receiving SONAR, optics or track manager auditory information, or inputting a solution). Data were epoched at each electrode from -1000ms before trigger onset to 1000ms after trigger onset. Grand average ERP visualisations were calculated from individual participant averages across the 2000ms window. Visual inspection of the ERP waveforms demonstrated a clear accumulation of negative amplitudes prior to participants’ setting a solution. For further analysis of this component, solution trials were epoched from -2000ms before trigger onset to 1000ms after trigger onset and individual participant averages for the amplitude at each electrode were created in windows from -1000ms to 0ms (at behavioural onset; setting a solution). No traditional baseline correction was applied as trial-by-trial mean ‘pre-stimulus’ voltage (in this case, from -1300ms to -1000ms) was included as a covariate in statistical analyses. This technique has been recommended by Alday (2019) to better consider the differences between conditions at baseline and increase statistical power through a reduction in the amount of variance in the residual error term.

### 3.4 Statistical Analysis

All statistical analyses and data visualisations were performed in *R* v4.3.2 and R Studio (R Core Team, 2020). The following packages were used: *tidyverse* v2.0.0 (Wickham et al., 2019), *ggeffects* v1.3.4 (Lüdecke, 2018), *lmerTest* v3.1.3 (Kuznetsova et al., 2017), *performance* v0.10.8 (Lüdecke et al., 2021), *car* v3.1.2 (Fox & Weisberg, 2019), *lme4* v1.1.35 (Bates et al., 2015), *ggpubr* 0.6.0 (Kassambra, 2023) and ggplot2 v3.4.4 (Wickham, 2016). Statistical analyses were run using *lme4*, *lmerTest* and *car*. To extract and visualise modelled effects, as well as descriptive data, *ggplot*, *ggeffects* and *ggpubr* were used. To create model output tables, *lmerOut* v0.5.1 (Alday, 2018) was used. Model fits were inspected using the *performance* package.

We used linear mixed-effects models (fit by restricted maximum likelihood estimates) to evaluate the predictive capacity of the variables of interest. To assess how adherence to the cognitive training regime related to on-task neural dynamics, we ran multiple models following a similar general formula:

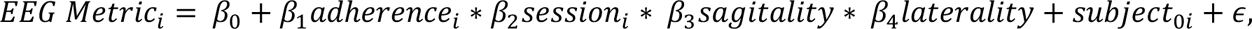

Here, the dependent variable (*EEG metric)* refers to the EEG variable of interest (alpha power, theta power, ERP amplitude, 1/*f* slope or 1/*f* intercept) summarised across all experimental epochs. Asterisks indicate interaction terms, including all subordinate main effects, while pluses denote additive terms. *Adherence* indicates the total number of cognitive training sessions completed (from 0 – 21 sessions), while *Sagitality* and *Laterality* express categorical variables relating to topographical electrode position. Electrodes were assigned to categories as follows: for the right laterality, anterior contained electrodes F4, F8, FC2 and FC6, central contained T8, C4, CP2 and CP6 while posterior contained P4, O2 and P8. For the left laterality, anterior contained electrodes F3, F7, FC1, and FC5, central contained T7, C3, CP1 and CP5, while posterior contained P3, O1 and P7. Midline electrodes included Fz, Cz and Pz and Oz. An additional control term, *pre-stimulus amplitude,* was included in models predicting ERP amplitude (as discussed above; Alday, 2019). Model intercepts and slopes were grouped by subject ID across session. Type II Wald tests from the *car* package were used to estimate *p-*values for each effect. Categorical factors were sum-to-zero contrast coded (whereby factor estimates are compared to the grand-mean; Schad et al., 2020). This technique allows the directionality of effects to be determined through visualisation of confidence intervals, circumventing the need for additional post-hoc testing (see Brehm & Alday, 2022 for a detailed discussion). Here, non-overlapping confidence intervals (at the 83% level) indicate significant differences at *p* < 0.05 (Austin & Hux, 2002; MacGregor-Fors & Payton, 2013).

## 4. Results

### 4.1 Descriptive Statistics

Detailed TMA task performance data is reported in Dziego, Bornkessel-Schlesewsky, Schlesewsky et al. (2024). On average, participants engaged in the mindfulness training sessions 9.8 times across the one-week period (SD = 6.98, range = 0 – 21). Here, the mean number of sessions per day was 1.47, with a linear increase in the average number of sessions per day (ß = 0.045, *p* = 0.278). By-participant adherence trends can be found within the supplementary materials (S4.1).

Average fluctuations across participants’ alpha power, theta power, and 1/*f* estimates across experimental time in both session 1 (pre-training) and session 2 (post-training) are shown in Figure 1. Mean estimates exhibited little fluctuation across experimental time. On average, participants demonstrated flatter slopes, lower intercepts, and decreased alpha and theta power during post-training (session 2) when compared to pre-training testing (session 1). Relationships between all variables of interest are depicted in Figure 2 (including individual resting-state metrics estimated in Dziego, Bornkessel-Schlesewsky, Schlesewsky et al., 2024). Here, we observed significant correlations between on-task alpha and theta power (R = -0.52, *p* < 0.001) and resting-state 1/*f* slopes and on-task slopes (R = 0.54, *p* < 0.001). On-task alpha and theta power were negatively related, whereby increased alpha was associated with decreased theta power. Moreover, mean TPE scores were significantly correlated with on-task theta power (R = 0.53, *p* < 0.001), in which higher theta power was related to larger TPE scores (signifying poorer performance). Average power and aperiodic estimates for each electrode across sessions are represented in Figure 3.

**Figure 1.**
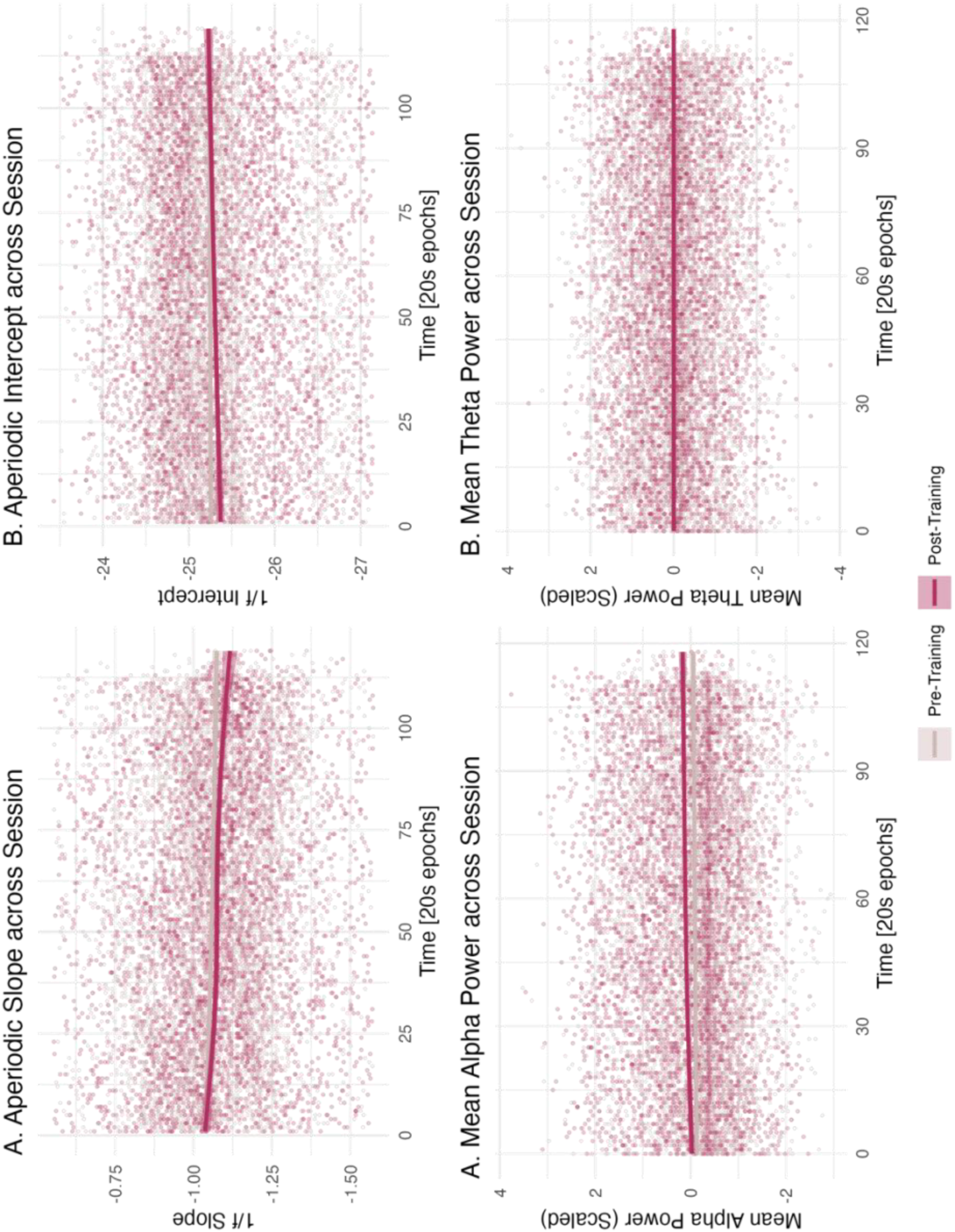
Average EEG metrics of interest across session 1 (pre-training) and session 2 (post-training). Aperiodic slope (A) and intercept (B) are depicted across the top row, while mean alpha power (C) and mean theta power (D) are depicted in the bottom row. The x-axis represents experimental time (in 20 second epochs). The y-axis represents the values of the EEG metric of interest. Pre-training values are shown in grey, while post-training values are shown in maroon. Individual points represent individual participant data.

**Figure 2.**
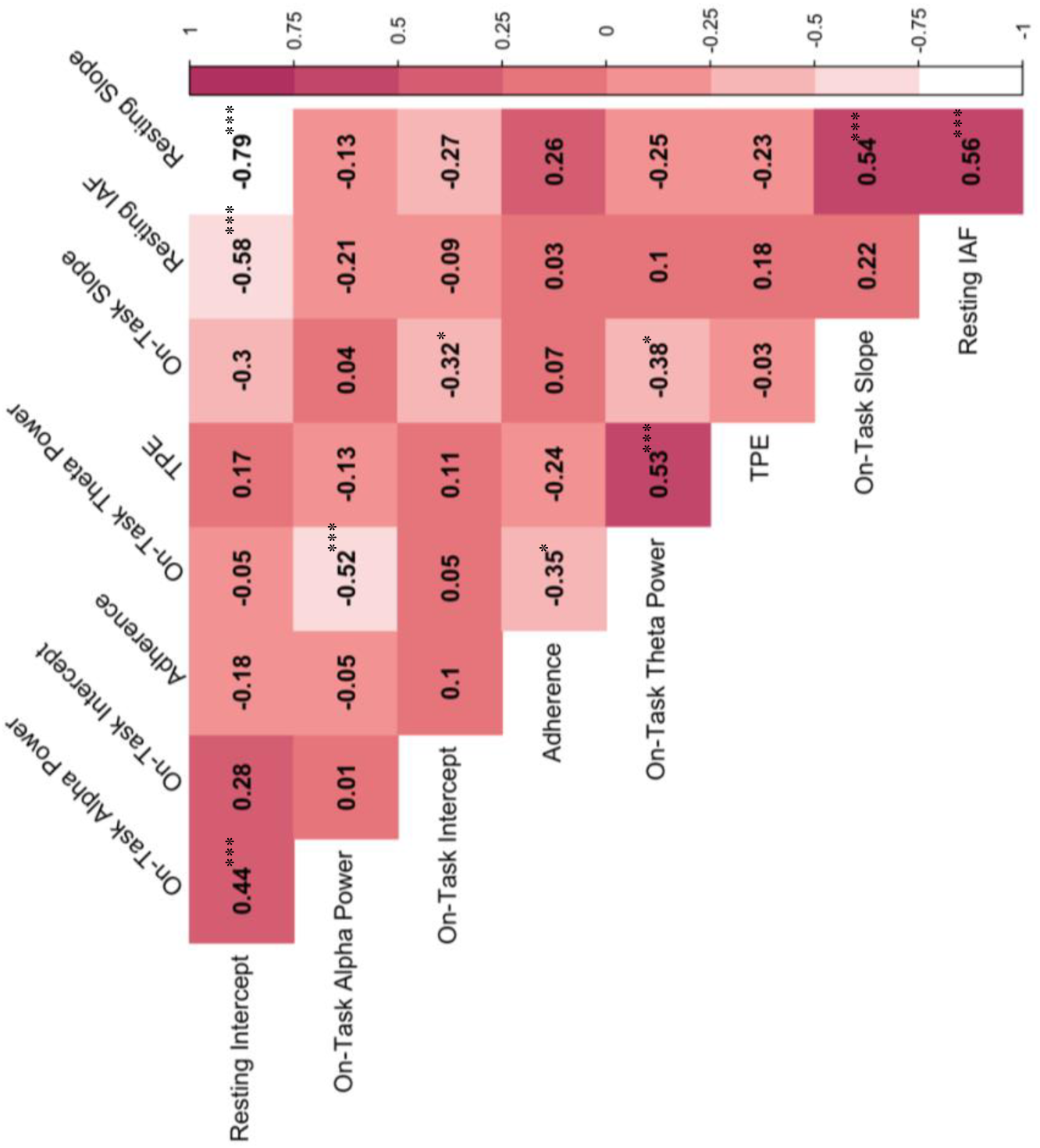
Relationship between adherence, mean TPE and on-task and resting-state EEG dynamics of interest (from session 1). Here, IAF denotes individual alpha frequency (calculated for individualised frequency bands) estimated during resting-state. Adherence denotes total sessions of cognitive training intervention completed. Correlogram illustrates the correlation coefficients between each of the metrics of interest. Negative (white) values indicate a negative association, while positive (maroon) values indicate a positive association. Values denote Pearson’s r coefficients. Note that statistically significant associations are denoted with a * (*p* < 0.5), ** (p < 0.01) and *** (*p* < 0.001). Plot created using the *corrplot* package.

**Figure 3.**
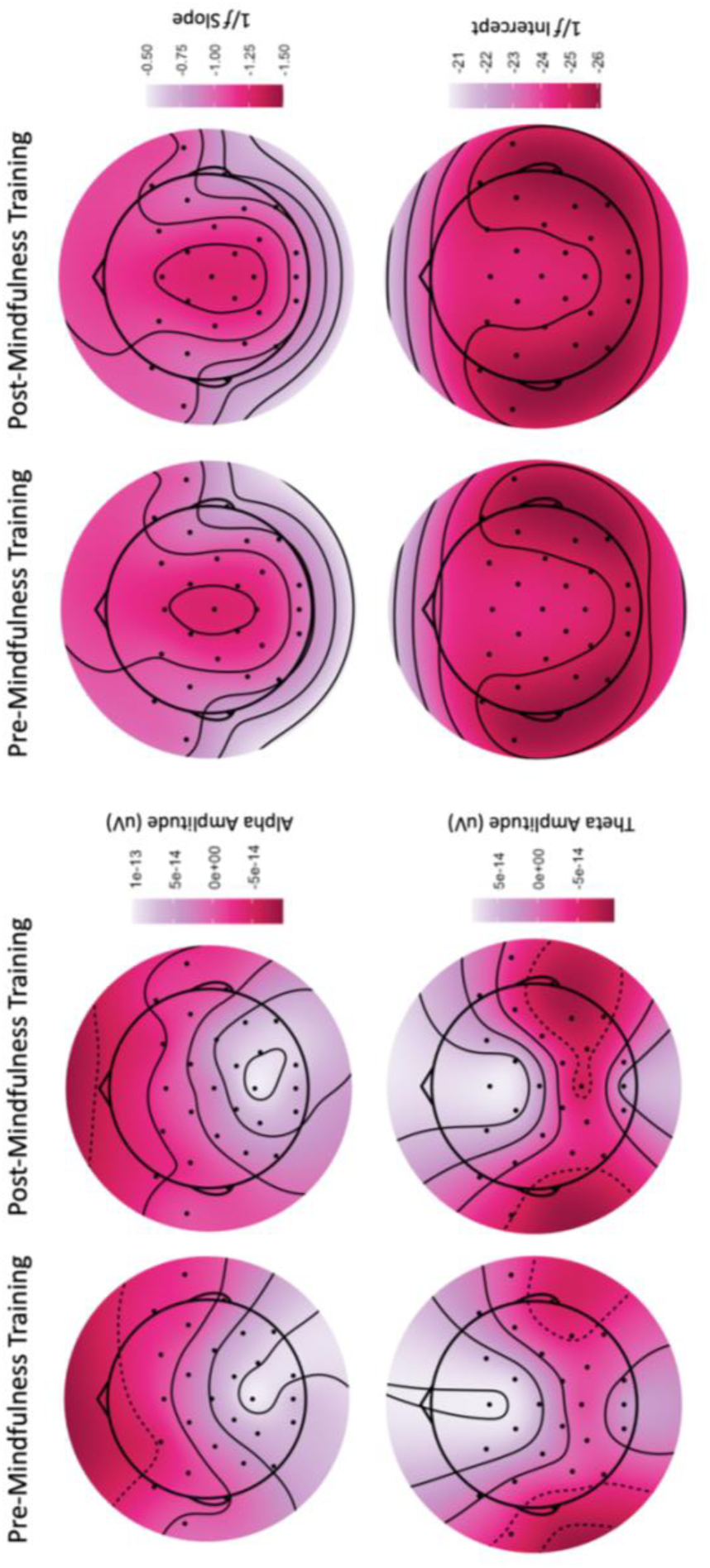
Topoplots demonstrating average alpha power, theta power, 1/*f* slopes and 1/*f* intercept during the TMA task (pre- and post-mindfulness training). Changes in alpha and theta power are demonstrated on the left, while changes in aperiodic 1/*f* activity are depicted on the right. Dark red indicates lower values, while lighter pink indicates higher values. Plots created using the *eegUtils* (Craddock, 2022) package in R.

### 4.2 Modelling On-Task Oscillatory and Aperiodic EEG Dynamics

To investigate how adherence to the cognitive training regime was associated with on-task EEG dynamics, we ran linear mixed-effects models predicting EEG activity from adherence levels, across session. All model outputs are depicted in supplementary materials (S4.2). For alpha power, we did not observe a significant interaction of *Adherence* x *Session* (X^2^(1) = 1.08, *p* = 0.298), demonstrating that alpha power did not vary across session by levels of adherence. However, a significant main effect of *Session* (X^2^(1) = 5.32, *p* < 0.05) demonstrated that alpha power was higher in session 2 (when compared to session 1) across all levels of adherence. Modelling of theta power demonstrated a significant interaction of *Sagitality* x *Laterality* (X^2^(4) = 44.99, *p* < 0.001), whereby theta power was higher overall in mid-anterior and mid-central electrodes. A significant main effect of *Adherence* (X^2^(1) = 4.56, *p* = 0.033) demonstrated increased theta power across all sessions (1 and 2) was also associated with individuals who adhered less to the cognitive training program.

For the 1/*f* slope, we did not observe a significant *Adherence* x *Session* (X^2^(1) = 0.92, *p* = 0.34) interaction, demonstrating levels of adherence did not significantly predict 1/*f* slopes during task engagement. A significant interaction of *Sagitality* x *Laterality* (X^2^(4) =18.87, *p* < 0.001) revealed the slope was steepest in middle electrodes, and flatter more posteriorly. A significant *Adherence* x *Sagitality* interaction (X^2^(1) = 20.43, *p* < 0.001) also demonstrated that higher levels of adherence were related to flatter slopes (in both session 1 and 2) in posterior regions (see Figure 4C). For the 1/*f* intercept, we observed a significant *Adherence* x *Session* x *Sagitality* interaction (X^2^(2) = 12.2, *p* = 0.002), whereby levels of adherence were associated with varying intercepts. Here, the largest difference between adherence was observed at session 1 in anterior electrodes (whereby those with less adherence demonstrated lower intercepts). For session 2, intercept values were more similar across *Sagitality*. A significant interaction of *Sagitality* x *Laterality* (X^2^(4) = 24.27, *p* < 0.001) demonstrated that intercept values were higher in mid-central and mid-anterior electrodes. In addition to this, significant interactions of *Adherence* x *Sagitality* (X^2^(2) = 6.31, *p* = 043) and *Adherence* x *Laterality* (X^2^(2) = 10.13, *p* = 0.006) demonstrated lower levels of adherence were related to more varied intercepts in left and right regions, while centre electrodes were more stable. Modelled effects across all EEG variables of interest are depicted in Figure 4.

**Figure 4.**
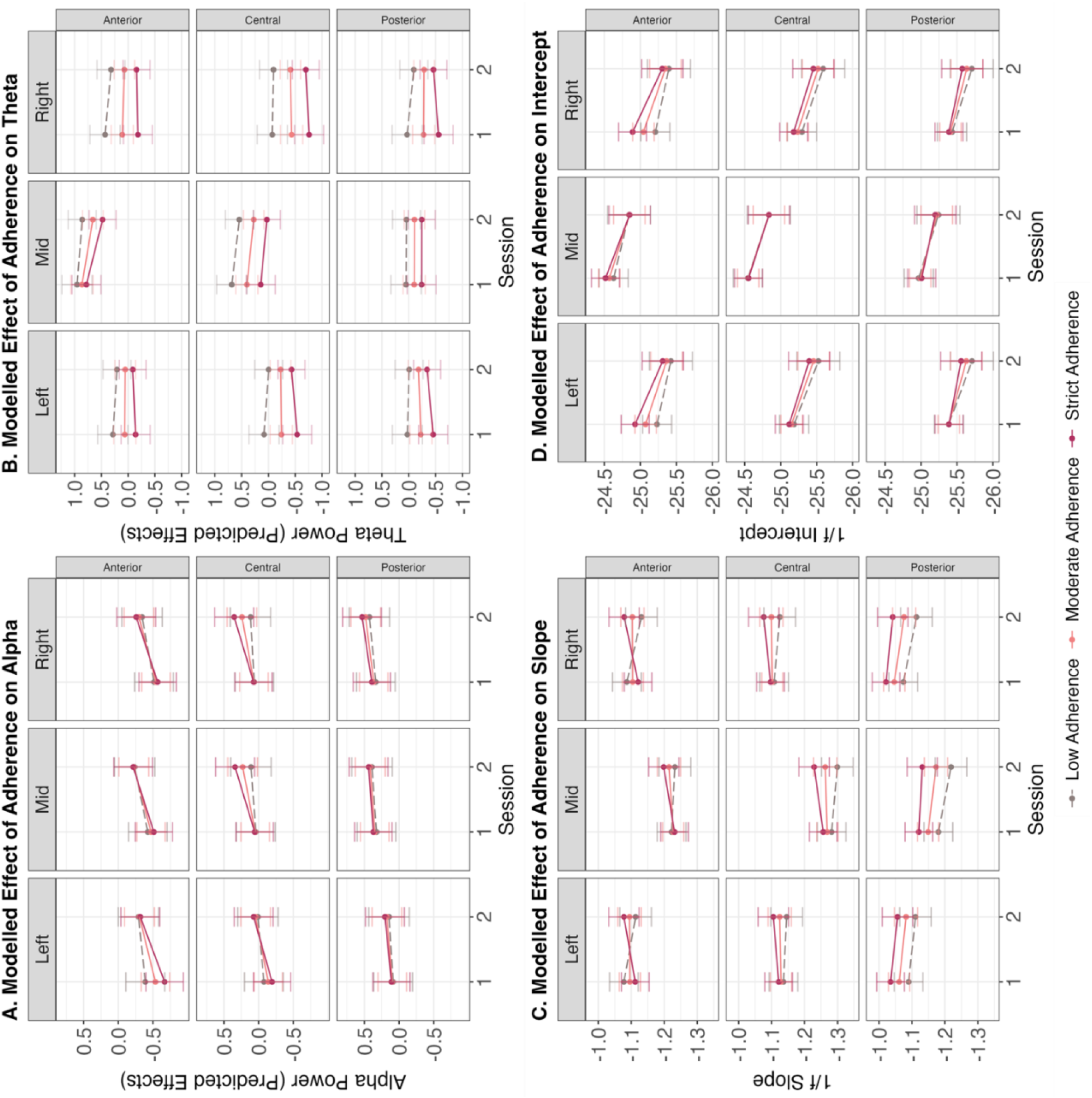
Modelled effects of adherence predicting EEG variables. Here, (A) depicts the relationship between alpha power and adherence, (B) demonstrated the relationship between theta power and adherence, while (C) demonstrates the modelled effects of adherence on 1/*f* slope and (D) for 1/*f* intercept. Colour represents levels of adherence to the cognitive training regime (ranging from 0-21) with lighter grey shades depicting poorer (low) adherence, and darker (maroon) shades for higher (strict) adherence. Session is shown on the x-axis.

### 4.3 Analysis of Event-Related Potentials

On average, each participants’ EEG recording included a total of 216.76 trials (SD = 71.27, range = 57 – 436) across all experimental triggers. This included an average of 56.87 trials for setting a solution (SD = 26.12, range = 13 – 154) and an average of 159.89 epochs around receiving auditory information (SD = 71.3, range = 57 – 436). This comprised a mean of 62.89 SONAR trials (SD = 21.1, range 17 – 145), 46.9 optics trials (SD = 14.61, range = 10 – 85) and 50.1 track manager trials (SD = 17.35, range = 13 – 154) across all participants. Grand average ERPs at Cz, grouped by adherence level (high = 15+ sessions, average = 4 – 14 sessions, low = <4 sessions) are shown in Figure 5. Visual inspection of the ERP plots demonstrated large variability in the amplitudes across time, particularly in the plots relating to auditory information. Here, grand average plots are lacking many of the clearly identifiable components related to stimulus presentation (N1/P2) and thus, no further inferential tests were conducted.

**Figure 5.**
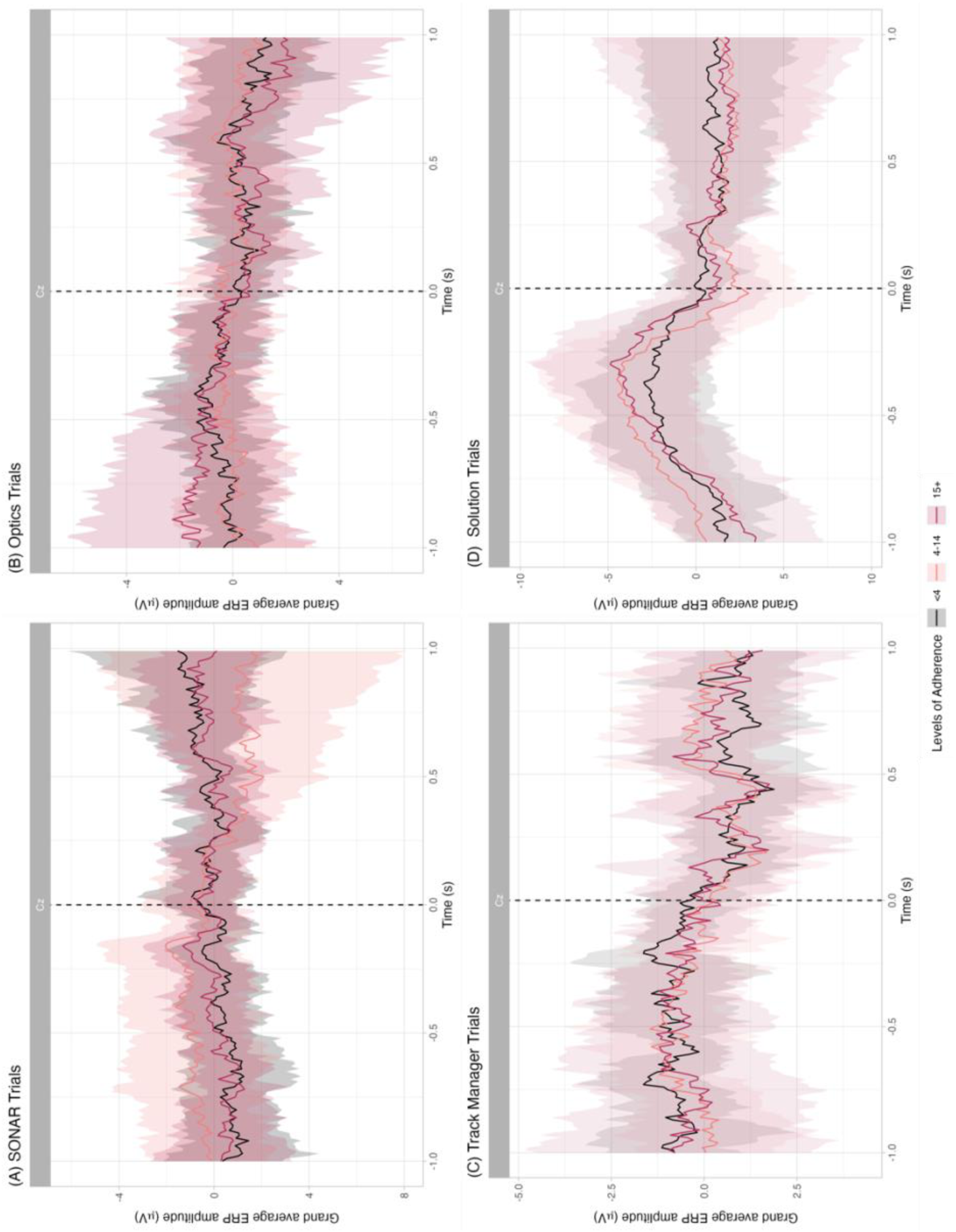
Grand average ERP waveforms (Cz) to all TMA task triggers. Here, grand averages depict waveforms observed when receiving auditory information from (A) SONAR, (B) optics and (C) the track manager and when setting a solution (D). Waveforms are time-locked (-1000ms to 1000ms) either to the onset of the auditory stream, or at participants’ click of the mouse (when inputting a solution). Darker red lines indicate higher adherence to the cognitive training regime (15+ sessions completed), pink indicates average adherence (between 4-14 sessions completed) and black indicates poorer adherence (<4 sessions completed). Negativity is plotted upwards. Surrounding fill (ribbons) represent +/-1 SD.

However, we note a clear cumulation of negative amplitudes prior to ‘setting a solution’, demonstrated in Figure 4D. For this component (i.e., from -1000ms to 0ms prior to behavioural onset), linear mixed models demonstrated a significant interaction of *Adherence* x *Session* x *Laterality* x *Sagitality* (X^2^(4) = 3.65, *p* < 0.001), whereby amplitudes were more similar across session for those with lower adherence but decreased across session for those with higher adherence. This was consistent across most regions, except in frontal, right regions, whereby amplitude increased for those with higher levels of adherence (see Figure 6 for (A) grand averages across electrodes and (B) modelled effects).

**Figure 6.**
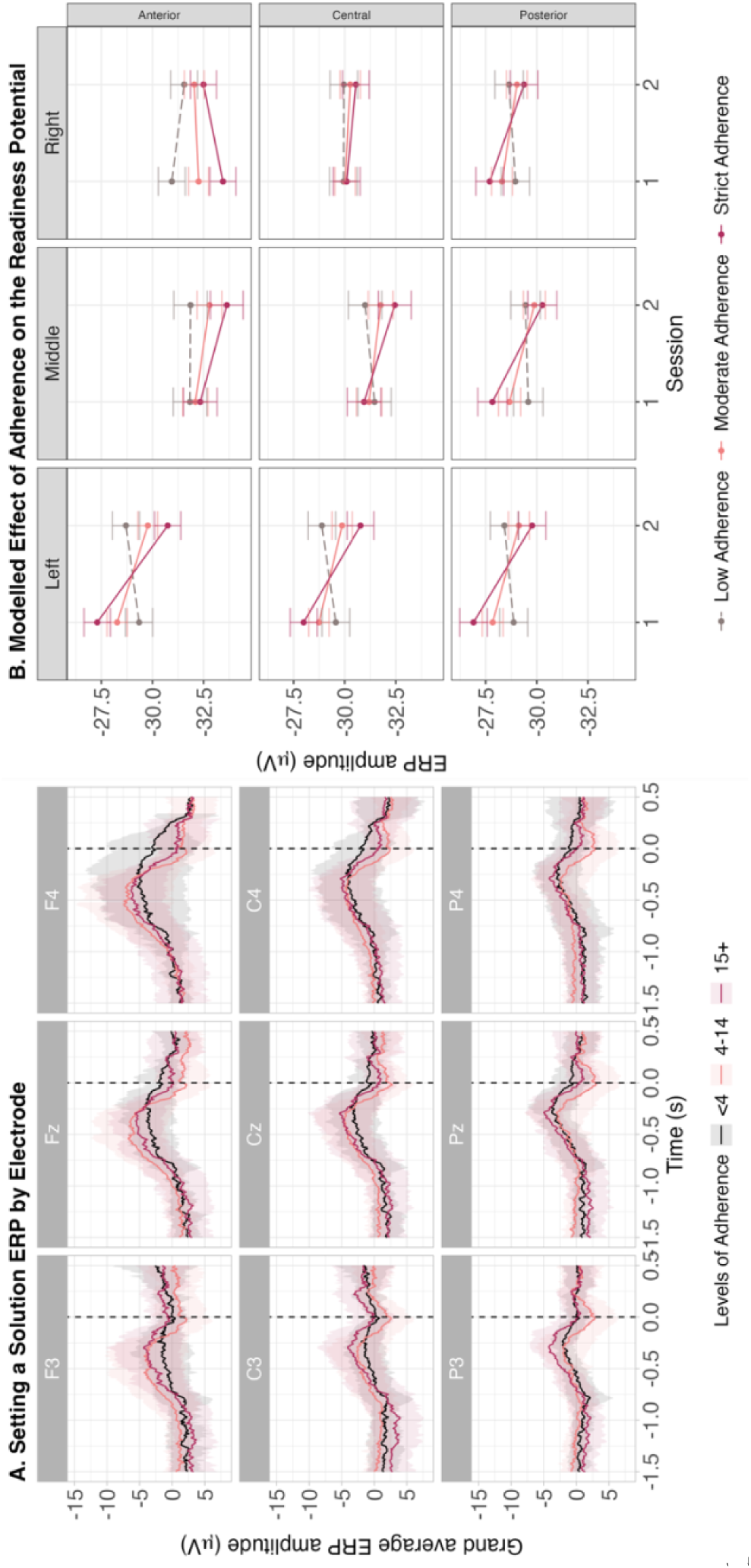
(A) Grand average ERP waveforms (by electrode) to setting a solution. Waveforms are time-locked (-2000ms to 1000ms) to participants’ click of the mouse. Darker red lines indicate higher adherence to the cognitive training regime (15+ sessions completed), pink indicates average adherence (between 4-14 sessions completed) and black indicates poorer adherence (<4 sessions completed). Negativity is plotted upwards. Surrounding fill (ribbons) represent +/-1 SD. (B) Modelled effects of adherence predicting amplitude. Colour represents levels of adherence to the cognitive training regime (ranging from 0-21) with lighter grey shades depicting poorer (low) adherence, and darker (maroon) shades for higher (strict) adherence. Session is shown on the x-axis.

## 5. Discussion

The current study aimed to explore EEG signatures related to cognitive enhancement from mindfulness-based cognitive training, specifically during cognitive task engagement. While studies have aimed to better understand *how* and *why* cognitive gains occur, current investigations have been limited by assessing EEG only during mindfulness engagement, comparing longer-term meditators/controls or focusing solely on ERPs, often overlooking the influence of aperiodic activity. Here, we explored a range of EEG components (oscillatory, aperiodic and event-related) to examine how mindfulness-based cognitive training adherence can be tracked during subsequent task engagement. Using linear mixed-effects models, we were unable to observe any clear predictive effect of levels of adherence across session on alpha power, theta power and 1/*f* slopes, opposing many of the previously observed relationships in mindfulness- and performance-related literature. We did, however, note topographical differences amongst alpha and theta power across cognitive task engagement and documented main effects of adherence or session, independently. Grand average ERPs around critical events (i.e., receiving SONAR information, or setting solutions) did not exhibit many of the established waveforms depicted in the mindfulness literature, highlighting the potential confounds when using event-related activity in this more complex paradigm. Nevertheless, we observed a lesser cumulation of negative amplitudes (prior to participants setting a solution) in session 2, in those who were high adherers.

### 5.1 Alpha and Theta Power

Our results demonstrated that levels of adherence to the mindfulness-based cognitive training regime did not predict differences in alpha power during the second testing session. These findings are inconsistent both with results reported *during* meditation (whereby alpha increases during engagement in meditation; e.g, Ahani et al., 2014; Cahn & Polich, 2006; Huang & Lo, 2009; Lagopoulos et al., 2009; Lomas et al., 2015; Takahashi et al., 2005) and in relation to superior cognitive performance (Bazanova & Vernon, 2014; Cooper et al., 2003; Klimesch, 1999; Klimesch et al., 2007), suggesting that the commonly observed neurophysiological signatures of meditation do not persist more longitudinally across other cognitive states. Further, as we were unable to detect any relationships between adherence in the training intervention and alpha power at session 2, our findings suggest that changes in alpha power dynamics do not directly parallel behavioural benefits (previously observed in Dziego, Bornkessel-Schlesewsky, Schlesewsky et al., 2024) garnered from mindfulness training. While these findings appear to contradict previous literature in the field (often associating mindfulness or enhanced cognition with increased alpha power), these results may point to a more nuanced relationship. For example, following mindful breathing exercises (Wong et al., 2018) or a longer 8-week intervention (Bing-Canar et al., 2016), improvements in the psychomotor vigilance or Stroop task are paralleled by changes in alpha desynchronisation. However, these studies specifically examined event-related desynchronisation (Pfurtscheller & Lopes da Silva, 1999), which is defined as the mean percentage of frequency synchronisation change (200-500ms post-trial onset). This approach largely differs to the methods presented within the current study but could suggest that a finer-grained analysis (e.g., focusing on oscillatory activity across smaller time scales) could provide more detailed insights.

Alternatively, while alpha power is considered to facilitate superior information processing by inhibiting irrelevant cortical areas (Bazanova & Vernon, 2014; Cooper et al., 2003; Klimesch et al., 2007), our findings could suggest that other cognitive processes may play a more critical role in the benefits gained from mindfulness training – at least in more complex environments (such as the TMA task). As previous analysis of this sample indicated that those with greater adherence exhibited superior performance (Dziego, Bornkessel-Schlesewsky, Schlesewsky et al., 2024), we would assume that alpha power would similarly demonstrate a clear relationship to levels of adherence. However, this was not the case, and average correlations between variables of interest (see Figure 2) demonstrated that mean alpha power was not associated with performance measures (i.e., TPE scores). Nevertheless, we observed a significant effect of session, demonstrating that alpha power was higher during the first testing session, when compared to second session, irrespective of levels of adherence (particularly evident in frontal and midline regions). This finding could be associated with the learning-related costs and differential allocation of attention during the first iteration of the TMA task, versus the second, where participants have a clearer understanding of their objectives. For example, alpha power attenuation has been shown to parallel task demands, whereby decreases are greater during more demanding cognitive tasks in EEG (Magosso et al., 2019) and MEG contexts (Wang et al., 2021, 2022), which might be the case during the first testing session of the TMA task.

In contrast to our hypotheses, we also did not observe a significant relationship between adherence, session and theta power. When considering the extant literature regarding EEG during meditation, these findings suggest that the commonly observed theta increases (e.g., Griffin et al., 2004; Klimesch, 1999; Seager et al., 2002) also do not transfer longitudinally (i.e., to other cognitive engagements/states) to explain enhancements in performance. However, it is critical here to note that we accounted for the 1/*f* slope within our power estimates (typically ignored by other researchers) which is known to have particularly powerful influence on theta estimates (Donoghue et al., 2020). In contrast to alpha power, we did not observe a significant change in theta power across the two testing sessions, regardless of levels of adherence. However, correlations between all variables of interest demonstrated that performance scores (TPE) were significantly correlated with on-task theta power, whereby higher theta power was associated with higher TPE scores (i.e., worse performance). As theta is associated with memory encoding and novel environments (Berry & Thompson, 1978; Klimesch et al., 1997; Seager et al., 2002), we might expect to see greater theta power in poorer performers (i.e., while participants are still ‘learning’ the task) and subsequent decreases with increased expertise.

Interestingly, we also observed a significant difference between levels of adherence and overall theta power, indicating a critical fundamental difference between our higher and lower intervention adherers. It is, however, difficult to report on how this finding relates to individuals who are higher or lower adherers. For example, elevated theta power during tasks has been linked to neuroticism and avoidance (Neo & McNaughton, 2011), motor impulsivity (i.e., “acting without thinking”; Leicht et al., 2013) or cognitive load (Chikhi et al., 2022; Puma et al., 2018). In the resting-state literature, enhanced theta power has been related to a multitude of different cognitive traits and capabilities (e.g., ADHD or changes with ageing; Finnigan & Robertson, 2011; Hermens et al., 2005; Tye et al., 2014) and is known to have a large genetic (stable) component (Tye et al., 2014; Zietsch et al., 2007). We could speculate on these links within our sample, but it is impossible to decipher what this may mean without comprehensive personality and demographic analyses.

Overall, our oscillatory analysis failed to demonstrate associations in on-task EEG dynamics as a product of adherence to mindfulness-based cognitive training. Our alpha and theta power findings add further uncertainty in the understanding of meditation- and performance-related EEG dynamics, suggesting that while these oscillations may serve as key signatures *during* meditation engagement, they may not be directly associated with adherence to mindfulness training (when measured during subsequent cognitive testing). While relationships between alpha and theta activity and performance have been demonstrated in more straightforward cognitive tasks (Bazanova & Vernon, 2014; Karakaş, 2020), these measures may be confounded by a variety of brain processes that are necessary during engagement in more dynamic environments. It is not atypical for previous findings in the field to not replicate in more naturalistic testing settings (Matusz et al., 2019; Virk et al., 2024). Alternatively, some researchers posit that cognitive changes from mindfulness-based interventions do not accumulate across longer durations (Lodha & Gupta, 2022), which might also be reflected in neurophysiology (e.g., with critical changes occurring in early time points, as measured through theta power; Lomas et al., 2015 and behavioural change; Dziego, Bornkessel-Schlesewsky, Schlesewsky et al., 2024). For example, it may only be one or two sessions that are necessary to evoke measurable neurophysiological change, which cannot be captured by our overall adherence-based analysis approach.

### 5.2 Aperiodic 1/*f* Change

Regarding aperiodic 1/*f* activity, we did not observe any relationship between adherence to the mindfulness-based cognitive training, session, and 1/*f* slopes during the TMA task. It is difficult to contextualise these findings amongst previous meditation and cognitive performance literature, as current explorations and knowledge on aperiodic activity are limited and still highly debated. This result does align with the findings within resting-state and complex task performance (in this same sample; Dziego, Bornkessel-Schlesewsky, Schlesewsky et al., 2024), whereby slopes did not change across the one-week intervention period. However, alternative research has demonstrated that changes to the 1/*f* slope from mindfulness training can persist more longitudinally, both during meditation (Rodriguez-Larios et al., 2021) and external to meditation engagement directly (Dziego, Zanesco, et al., 2024). In cognitive performance settings, investigations of the 1/*f* slope during task engagement are scarce, but Cross, Corcoran, et al. (2022), Sheehan et al., (2018; in intracranial EEG) and Pertermann et al. (2019) demonstrated that slopes track performance. For Cross, Corcoran et al. (2022) and Sheehan et al. (2018), flatter 1/*f* slopes were linked to superior performance outcomes; thought to be related to greater complexity in neural signals, providing a foundation for superior integration, flexibility, memory capacity and processing of information in more complex tasks (Dziego, Bornkessel-Schlesewsky, Schlesewsky et al., 2024; Medel et al., 2023; Tran et al., 2020). On top of this, results by Pertermann et al. (2019) indicated that the 1/*f* slope steepens where cognitive control and response inhibition is required, when compared to instances where responses need to be executed. The continual and persistent nature of the TMA task may foster a cognitive strategy focusing on consistent response execution. Nevertheless, it is difficult to rectify these results, as they stem from largely varying routes of inquiry and methodological stances. In a similar vein to our theta findings, we also observed that 1/*f* slopes were associated with levels of adherence (across both sessions 1 and 2), whereby slopes were flatter for higher adherers (versus low adherers), particularly in posterior regions. As above, this may demonstrate a fundamental difference (whether in personality, cognitive strategies, motivation or neural dynamics) between levels of adherence.

Assessing the 1/*f* parameters *during* task engagement (as opposed to during the resting-state) brings up a critical consideration within this area of research. As investigating aperiodic 1/*f* activity is relatively novel, there is still debate about the best way in which to utilise this metric. For example, some work aims to use this parameter as a trait-like marker (e.g., (Cross, Chatburn, et al., 2022; Dziego et al., 2023; Immink et al., 2021; Ouyang et al., 2020), whereby the slope and intercept are viewed as stable characteristics of an individual (i.e., like a biological fingerprint; Demuru & Fraschini, 2020; Pathania et al., 2021). Other studies have viewed 1/*f* activity as a more fluid, flexible, state-like marker (e.g., Chatburn et al., 2024; Colombo et al., 2019; Lendner et al., 2020), whereby the slope changes across arousal levels, sleepiness and states of consciousness. Here, we observed that the 1/*f* parameters generally exhibited little fluctuation across experimental time, despite dynamic changes in task difficulty, suggesting a more trait-like function (at least in the wakeful state). Moreover, we observed a small correlation between resting-state 1/*f* and on-task 1/*f* (see Figure 2), which suggests that while these two components are linked to some extent, there may be both trait-like and state-like fluctuations within this EEG parameter. While the literature is beginning to enquire further into these markers (e.g., Chatburn et al., 2023; Demuru & Fraschini, 2020; Lendner et al., 2020; Pathania et al., 2021; Schaworonkow & Voytek, 2021), more experimentation is required to clearly detail the intra- and inter-individual fluctuations in these parameters and how they relate to cognition, states of consciousness and development over time.

For the 1/*f* intercept, we did not observe significant adherence by session effects, but lower intercepts for those with less adherence were observed in the first testing session (in left and right frontal regions). It is more difficult to speculate what this may functionally and behaviourally indicate, as the research surrounding the 1/*f* intercept is also incredibly limited. However, the intercept has been related to overall neuronal population spiking (Manning et al., 2009; Miller et al., 2009) and global field power (Dziego, Zanesco, et al., 2024). From this perspective, the 1/*f* intercept appears to track overall neural activity across the frequency spectrum (Gardony et al., 2017; Jaušovec & Jaušovec, 2000; Mölle et al., 1996) but it is unknown what it may reveal about lower adherers in the first session.

### 5.3 Event-Related Potentials

As an additional exploratory investigation, we created grand average ERPs surrounding critical events in the TMA task (i.e., receiving auditory information and setting a solution). Here, we observed that (particularly for auditory information triggers), ERP waveforms were highly variable and lacking the meaningful waveforms commonly established in previous literature. Reflecting on the TMA task, we note that this more naturalistic simulation environment may be too ‘noisy’ for traditional smaller-scale ERP investigations. Classically, ERPs are computed from recordings during computer-based tasks, whereby participants exhibit minimal movement. The TMA task occurs across dual computer screens (encouraging consistent movement and scanning by participants) which may introduce considerable artefacts into the recordings. Nevertheless, studies have shown that established ERPs (like the mismatch negativity) can still be elicited under more naturalistic conditions (i.e., walking around university campus; Liebherr et al., 2021), suggesting that other confounds may play a larger role.

In this case, we note that trials were inconsistent across participants, with the number of triggers within each EEG recording dependent on the specific number of times participants clicked to receive auditory information or set solutions. Large discrepancies in the number of trials for each participant are likely impacting the equal influence of all separable participant trials, potentially skewing the grand average results. Lastly, we note that unlike traditional ERP paradigms, it is not known whether participants were truly engaging with the auditory information received. During the TMA task, multi-modal information is presented simultaneously at any given moment. Participants must be watching various modules on each screen (e.g., the geo-plot or dot stack) and inputting numeric parameters, all while audio and written information about contacts is offered. We note that participants can also request certain auditory information to be replayed, but this can only occur after existing audio material is presented in full. Often, this can mean that participants actively ignore irrelevant audio outputs, while waiting for new information to be played. Overall, the TMA task lacks the repetitive, controlled structure of traditional ERP investigations (i.e., consistent trials, stimulus presentation durations, and clear inter-stimulus intervals), demonstrating how trial-based analysis can be more difficult in naturalistic testing settings (Sonkusare et al., 2019).

Nevertheless, there was one condition (setting a solution) in which we were able to identify an ERP component in the grand averages most clearly. This component can be observed as a slow negative-going amplitude increasing in the lead up to an onset of a motor movement, typically referred to as the Readiness Potential (Schurger et al., 2021). It is posited to reflect the planning and preparation of action, signalling self-initiated movement (Libet et al., 1983; Schurger et al., 2021). In relation to the ERP studies explored in the introduction, the Readiness Potential demonstrates a likeness to the CNV and in some cases, researchers have argued that these components are strongly related (Rohrbaugh et al., 1976; Rohrbaugh & Gaillard, 1983; Schlegel et al., 2013).

The evocation of a motor-related Readiness Potential in this context conforms to expectations, in reflecting upon its occurrence prior to participants selecting the ‘set solution’ button during the TMA task. Mixed modelling specifically demonstrated that this component was related to levels of adherence across time, where amplitudes were increased in session 2 for those with lower levels of adherence. These results contradict previous research of meditation and the Readiness Potential, where early neural activity was associated with participants’ reports of initiating intentional action, but differences in this component were not observed when controls were compared with experienced meditators (Jo et al., 2014).

Theoretically, this relationship may be exhibited due to relative differences in the required accumulation of neural activity (which may be likened to evidence accumulation models; Boag et al., 2023) prior to the onset of action initiation (or ‘setting a solution’). In the case of the TMA task, it may suggest that those who adhered more to the cognitive training regime experienced a smaller accumulation threshold (i.e., actions are initiated with less neural activity) when compared to those who adhered to a lesser extent (Lui et al., 2021; Travers & Haggard, 2020). In this case, participants who adhered more to the mindfulness-based cognitive training may require less information or be more confident in their parameters in the decision-making process of setting a solution (reaching the ‘decision boundary’ more rapidly). On the other hand, those with lower adherence may need to expend more neural resources to gather and utilise the necessary information to appropriately set a solution. In this regard, adherence to mindfulness interventions may reduce mental expenditure during cognitive and motor tasks.

### 5.4 Limitations and Future Directions

The present study involved a comprehensive examination of multiple EEG-related phenomena during task engagement. Although such electrophysiology can be tied to cognitive enhancement from mindfulness training, there are a few limitations to the exploratory investigation. Firstly, our novel approach to using aperiodic activity (Donoghue et al., 2020) and individualised frequency bands (Klimesch, 2012) makes it difficult to generalise results to previous oscillatory findings in the field. It is unknown how the inclusion and consideration of aperiodic activity or individualised bands in previous studies may alter the results, and thus it is hard to speculate how previous findings relate to these new observations. Secondly, we focused only on two specific oscillatory bands (most relevant to performance; alpha and theta) but did not examine power across higher frequencies (i.e., beta, delta or gamma). While less explored in the context of cognitive performance, several studies have shown that gamma activity is associated with memory encoding, cognitive performance, and impaired information processing (Başar, 2013; Herrmann et al., 2010; Jaušovec & Jaušovec, 2005). Likewise, research has explored the theta/beta ratio as a critical indicator of information processing capabilities. For example, associations have been seen between this ratio and self-reported attentional capacity and are speculated to reflect suboptimal functioning of the prefrontal cortex (Putman et al., 2010, 2014). These alternative oscillatory frequencies could provide more effective avenues in which mindfulness-based cognitive training adherence could be traced.

We hypothesise that overall, more nuanced analysis procedures in future research could help to better uncover the underlying mechanisms (measured through EEG) of mindfulness-induced performance change. For example, we observed a negative correlation between mean on-task alpha and theta power, whereby as alpha power increased, theta power decreased. This suggests that exploring the inter-connected nature of alpha and theta power (as detailed by Klimesch, 1999) or alpha-theta cross-frequency dynamics (see Rodriguez-Larios et al., 2020) in this domain may provide clearer insights. Further, Bing-Canar et al. (2016) and Wong et al., (2018) used event-related desynchronisation to explore mindfulness and cognitive performance, which may better predict benefits gained. These relationships suggest that mindfulness may work to induce change on the brain from a more dynamical perspective (i.e., comprising of the inter-relationships of different frequencies or neural dynamics), as opposed to directly influencing more independent measures.

Additionally, our exploration of ERPs around critical events within the simulation (i.e., receiving auditory information) was not particularly effective in distinguishing those with high versus low levels of adherence. In the future, naturalistic paradigms could be better catered to ERP analysis by including a more nuanced triggering system (capturing multiple aspects of the cognitive task simultaneously) and understanding the influence of multi-faceted cognitive engagements. Nevertheless, our study provides a bridge between more traditional lab-experiments and more ecologically valid performance settings and highlights critical factors that need to be considered during this transition.

### 5.5 Conclusion

The current study examined how adherence to a short-term mindfulness-based cognitive training regime was related to EEG dynamics during subsequent task engagement. Our findings demonstrated that EEG dynamics during task engagement were not predicted by levels of adherence across time, with no intervention-related associations found between alpha power, theta power, and 1/*f* parameters (slope and intercept). We also used ERPs to investigate time-locked changes in amplitude around critical events within the simulation. However, the more naturalistic task setting produced indistinct ERP waveforms, unfit for further inferential analyses. We did, however, see a clear cumulation of negative amplitudes prior to one condition (i.e., behavioural onset), which significantly differed by levels of adherence. This was interpreted as a Readiness Potential, possibly signifying differences in neural ‘accumulation’ required to initiate action during the TMA task. In summary, the present findings suggest that higher-order cognitive enhancements from a short-term mindfulness-based training regime are not easily traced through certain EEG parameters (alpha power, theta power, 1/*f* activity), and alternative analysis strategies may be better suited to understanding this relationship. Our ERP investigation demonstrated the potential confounds in the transferability of ERPs to more naturalistic, real-world contexts, but suggested that some components may be better positioned to exhibit traceable changes in dynamic tasks than others.

## References

1. Activity at Rest and During Evoked Negative Emotions. International Journal of Neuroscience, 115(6), 893–909. 10.1080/00207450590897969

2. Ahani, A., Wahbeh, H., Nezamfar, H., Miller, M., Erdogmus, D., & Oken, B. (2014). Quantitative change of EEG and respiration signals during mindfulness meditation. Journal of NeuroEngineering and Rehabilitation, 11(1), 87. 10.1186/1743-0003-11-87

3. Ainsworth, B., Eddershaw, R., Meron, D., Baldwin, D. S., & Garner, M. (2013). The effect of focused attention and open monitoring meditation on attention network function in healthy volunteers. Psychiatry Research, 210(3), 1226–1231. 10.1016/j.psychres.2013.09.002

4. Alday, P. M. (2018). lmerOut: LaTeX Output for Mixed Effects Models with lme4. (0.5) [Computer software]. https://bitbucket.org/palday/lmerout

5. Alday, P. M. (2019). How much baseline correction do we need in ERP research? Extended GLM model can replace baseline correction while lifting its limits. Psychophysiology, 56(12). 10.1111/psyp.13451

6. Amihai, I., & Kozhevnikov, M. (2014). Arousal vs. Relaxation: A Comparison of the Neurophysiological and Cognitive Correlates of Vajrayana and Theravada Meditative Practices. PLOS ONE, 9(7), e102990. 10.1371/journal.pone.0102990

7. Austin, P. C., & Hux, J. E. (2002). A brief note on overlapping confidence intervals. Journal of Vascular Surgery, 36(1), 194–195. 10.1067/mva.2002.125015

8. Başar, E. (2013). A review of gamma oscillations in healthy subjects and in cognitive impairment. International Journal of Psychophysiology, 90(2), 99–117. 10.1016/j.ijpsycho.2013.07.005

9. Bates, D., Mächler, M., Bolker, B., & Walker, S. (2015). Fitting Linear Mixed-Effects Models Using lme4. Journal of Statistical Software, 67(1), Article 1. 10.18637/jss.v067.i01

10. Bazanova, O. M., & Vernon, D. (2014). Interpreting EEG alpha activity. Neuroscience & Biobehavioral Reviews, 44, 94–110. 10.1016/j.neubiorev.2013.05.007

11. Benedek, M., Schickel, R. J., Jauk, E., Fink, A., & Neubauer, A. C. (2014). Alpha power increases in right parietal cortex reflects focused internal attention. Neuropsychologia, 56, 393–400. 10.1016/j.neuropsychologia.2014.02.010

12. Berry, S. D., & Thompson, R. F. (1978). Prediction of learning rate from the hippocampal electroencephalogram. *Science (New York*, N.Y.), 200(4347), 1298–1300. 10.1126/science.663612

13. Bing-Canar, H., Pizzuto, J., & Compton, R. J. (2016). Mindfulness-of-breathing exercise modulates EEG alpha activity during cognitive performance. Psychophysiology, 53(9), 1366–1376. 10.1111/psyp.12678

14. Boag, R. J., Strickland, L., Heathcote, A., Neal, A., Palada, H., & Loft, S. (2023). Evidence accumulation modelling in the wild: Understanding safety-critical decisions. Trends in Cognitive Sciences, 27(2), 175–188. 10.1016/j.tics.2022.11.009

15. Brehm, L., & Alday, P. M. (2022). Contrast coding choices in a decade of mixed models. Journal of Memory and Language, 125, 104334. 10.1016/j.jml.2022.104334

16. Britton, W. B., Lepp, N. E., Niles, H. F., Rocha, T., Fisher, N. E., & Gold, J. S. (2014). A randomized controlled pilot trial of classroom-based mindfulness meditation compared to an active control condition in sixth-grade children. Journal of School Psychology, 52(3), 263–278. 10.1016/j.jsp.2014.03.002

17. Brown, J. O., Chatburn, A., Wright, D. L., & Immink, M. A. (2022). A Single Session of Mindfulness Meditation Expedites Immediate Motor Memory Consolidation to Improve Wakeful Offline Learning. Journal of Motor Learning and Development, 11(1), 45–70. 10.1123/jmld.2022-0016

18. Cahn, B. R., Delorme, A., & Polich, J. (2010). Occipital gamma activation during Vipassana meditation. Cognitive Processing, 11(1), 39–56. 10.1007/s10339-009-0352-1

19. Cahn, B. R., & Polich, J. (2006). Meditation states and traits: EEG, ERP, and neuroimaging studies. Psychological Bulletin, 132(2), 180–211. 10.1037/0033-2909.132.2.180

20. Cahn, B. R., & Polich, J. (2009). Meditation (Vipassana) and the P3a Event-Related Brain Potential. International Journal of Psychophysiology : Official Journal of the International Organization of Psychophysiology, 72(1), 51–60. 10.1016/j.ijpsycho.2008.03.013

21. Chatburn, A., Lushington, K., & Cross, Z. R. (2024). Considerations towards a neurobiologically-informed EEG measurement of sleepiness. Brain Research, 1841, 149088. 10.1016/j.brainres.2024.149088

22. Chikhi, S., Matton, N., & Blanchet, S. (2022). EEG power spectral measures of cognitive workload: A meta-analysis. Psychophysiology, 59(6), e14009. 10.1111/psyp.14009

23. Colombo, M. A., Napolitani, M., Boly, M., Gosseries, O., Casarotto, S., Rosanova, M., Brichant, J.-F., Boveroux, P., Rex, S., Laureys, S., Massimini, M., Chieregato, A., & Sarasso, S. (2019). The spectral exponent of the resting EEG indexes the presence of consciousness during unresponsiveness induced by propofol, xenon, and ketamine. NeuroImage, 189, 631–644. 10.1016/j.neuroimage.2019.01.024

24. Compton, R. J., Gearinger, D., & Wild, H. (2019). The wandering mind oscillates: EEG alpha power is enhanced during moments of mind-wandering. *Cognitive, Affective*, & Behavioral Neuroscience, 19(5), 1184–1191. 10.3758/s13415-019-00745-9

25. Cooper, N. R., Croft, R. J., Dominey, S. J. J., Burgess, A. P., & Gruzelier, J. H. (2003). Paradox lost? Exploring the role of alpha oscillations during externally vs. internally directed attention and the implications for idling and inhibition hypotheses. International Journal of Psychophysiology: Official Journal of the International Organization of Psychophysiology, 47(1), 65–74. 10.1016/s0167-8760(02)00107-1

26. Corby, J. C., Roth, W. T., Zarcone, V. P., & Kopell, B. S. (1978). Psychophysiological correlates of the practice of Tantric Yoga meditation. Archives of General Psychiatry, 35(5), 571–577. 10.1001/archpsyc.1978.01770290053005

27. Craddock, M. (2022). eegUtils: Utilities for Electroencephalographic (EEG) Analysis. [Computer software].

28. Crane, R. S., Brewer, J., Feldman, C., Kabat-Zinn, J., Santorelli, S., Williams, J. M. G., & Kuyken, W. (2017). What defines mindfulness-based programs? The warp and the weft. Psychological Medicine, 47(6), 990–999. 10.1017/S0033291716003317

29. Cross, Z. R., Chatburn, A., Melberzs, L., Temby, P., Pomeroy, D., Schlesewsky, M., & Bornkessel-Schlesewsky, I. (2022). Task-related, intrinsic oscillatory and aperiodic neural activity predict performance in naturalistic team-based training scenarios. Scientific Reports, 12(1), Article 1. 10.1038/s41598-022-20704-8

30. Cross, Z. R., Corcoran, A. W., Schlesewsky, M., Kohler, M. J., & Bornkessel-Schlesewsky, I. (2022). Oscillatory and Aperiodic Neural Activity Jointly Predict Language Learning. Journal of Cognitive Neuroscience, 34(9), 1630–1649. 10.1162/jocn_a_01878

31. Dave, S., Brothers, T. A., & Swaab, T. Y. (2018). 1/f Neural Noise and Electrophysiological Indices of Contextual Prediction in Aging. Brain Research, 1691, 34–43. 10.1016/j.brainres.2018.04.007

32. Delgado-Pastor, L. C., Perakakis, P., Subramanya, P., Telles, S., & Vila, J. (2013). Mindfulness (Vipassana) meditation: Effects on P3b event-related potential and heart rate variability. International Journal of Psychophysiology, 90(2), 207–214. 10.1016/j.ijpsycho.2013.07.006

33. Demuru, M., & Fraschini, M. (2020). EEG fingerprinting: Subject-specific signature based on the aperiodic component of power spectrum. Computers in Biology and Medicine, 120, 103748. 10.1016/j.compbiomed.2020.103748

34. Dentico, D., Bachhuber, D., Riedner, B. A., Ferrarelli, F., Tononi, G., Davidson, R. J., & Lutz, A. (2018). Acute effects of meditation training on the waking and sleeping brain: Is it all about homeostasis? European Journal of Neuroscience, 48(6), 2310–2321. 10.1111/ejn.14131

35. Dietl, T., Dirlich, G., Vogl, L., Lechner, C., & Strian, F. (1999). Orienting response and frontal midline theta activity: A somatosensory spectral perturbation study. Clinical Neurophysiology: Official Journal of the International Federation of Clinical Neurophysiology, 110(7), 1204–1209. 10.1016/s1388-2457(99)00057-7

36. Donoghue, T., Haller, M., Peterson, E. J., Varma, P., Sebastian, P., Gao, R., Noto, T., Lara, A. H., Wallis, J. D., Knight, R. T., Shestyuk, A., & Voytek, B. (2020). Parameterizing neural power spectra into periodic and aperiodic components. Nature Neuroscience, 23(12), Article 12. 10.1038/s41593-020-00744-x

37. Dziego, C. A., Bornkessel-Schlesewsky, I., Jano, S., Chatburn, A., Schlesewsky, M., Immink, M. A., Sinha, R., Irons, J., Schmitt, M., Chen, S., & Cross, Z. R. (2023). Neural and cognitive correlates of performance in dynamic multi-modal settings. Neuropsychologia, 180, 108483. 10.1016/j.neuropsychologia.2023.108483

38. Dziego, C. A., Bornkessel-Schlesewsky, I., Schlesewsky, M., Sinha, R., Immink, M. A., & Cross, Z. R. (2024). Augmenting complex and dynamic performance through mindfulness-based cognitive training: An evaluation of training adherence, trait mindfulness, personality and resting-state EEG. PloS One, 19(5), e0292501. 10.1371/journal.pone.0292501

39. Edwards, J., Peres, J., Monti, D. A., & Newberg, A. B. (2012). The neurobiological correlates of meditation and mindfulness. In Exploring frontiers of the mind–brain relationship (pp. 97–112). Springer Science + Business Media. 10.1007/978-1-4614-0647-1_6

40. Elias, A. N., Guich, S., & Wilson, A. F. (2000). Ketosis with enhanced GABAergic tone promotes physiological changes in transcendental meditation. Medical Hypotheses, 54(4), 660–662. 10.1054/mehy.1999.0921

41. Fell, J., Axmacher, N., & Haupt, S. (2010). From alpha to gamma: Electrophysiological correlates of meditation-related states of consciousness. Medical Hypotheses, 75(2), 218–224. 10.1016/j.mehy.2010.02.025

42. Finnigan, S., & Robertson, I. H. (2011). Resting EEG theta power correlates with cognitive performance in healthy older adults. Psychophysiology, 48(8), 1083–1087. 10.1111/j.1469-8986.2010.01173.x

43. Fox, J., & Weisberg, S. (2019). An R Companion to Applied Regression. SAGE Publications.

44. Gallant, S. N. (2016). Mindfulness meditation practice and executive functioning: Breaking down the benefit. Consciousness and Cognition, 40, 116–130. 10.1016/j.concog.2016.01.005

45. Gardony, A. L., Eddy, M. D., Brunyé, T. T., & Taylor, H. A. (2017). Cognitive strategies in the mental rotation task revealed by EEG spectral power. Brain and Cognition, 118, 1–18. 10.1016/j.bandc.2017.07.003

46. Goldman, R. I., Stern, J. M., Engel, J., & Cohen, M. S. (2002). Simultaneous EEG and fMRI of the alpha rhythm. Neuroreport, 13(18), 2487–2492. 10.1097/01.wnr.0000047685.08940.d0

47. Griffin, A. L., Asaka, Y., Darling, R. D., & Berry, S. D. (2004). Theta-Contingent Trial Presentation Accelerates Learning Rate and Enhances Hippocampal Plasticity During Trace Eyeblink Conditioning. Behavioral Neuroscience, 118(2), 403–411. 10.1037/0735-7044.118.2.403

48. He, B. J. (2014). Scale-free brain activity: Past, present, and future. Trends in Cognitive Sciences, 18(9), 480–487. 10.1016/j.tics.2014.04.003

49. Hermens, D. F., Soei, E. X. C., Clarke, S. D., Kohn, M. R., Gordon, E., & Williams, L. M. (2005). Resting EEG theta activity predicts cognitive performance in attention-deficit hyperactivity disorder. Pediatric Neurology, 32(4), 248–256. 10.1016/j.pediatrneurol.2004.11.009

50. Herrmann, C. S., Fründ, I., & Lenz, D. (2010). Human gamma-band activity: A review on cognitive and behavioral correlates and network models. Neuroscience & Biobehavioral Reviews, 34(7), 981–992. 10.1016/j.neubiorev.2009.09.001

51. Herweg, N. A., Solomon, E. A., & Kahana, M. J. (2020). Theta Oscillations in Human Memory. Trends in Cognitive Sciences, 24(3), 208–227. 10.1016/j.tics.2019.12.006

52. Huang, H.-Y., & Lo, P.-C. (2009). EEG dynamics of experienced Zen meditation practitioners probed by complexity index and spectral measure. Journal of Medical Engineering & Technology, 33(4), 314–321. 10.1080/03091900802602677

53. Immink, M. A., Cross, Z. R., Chatburn, A., Baumeister, J., Schlesewsky, M., & Bornkessel-Schlesewsky, I. (2021). Resting-state aperiodic neural dynamics predict individual differences in visuomotor performance and learning. bioRxiv, 2021.04.08.438941. 10.1101/2021.04.08.438941

54. Incagli, F., Tarantino, V., Crescentini, C., & Vallesi, A. (2020). The Effects of 8-Week Mindfulness-Based Stress Reduction Program on Cognitive Control: An EEG Study. Mindfulness, 11(3), 756–770. 10.1007/s12671-019-01288-3

55. Jaušovec, N., & Jaušovec, K. (2000). Correlations between ERP parameters and intelligence: A reconsideration. Biological Psychology, 55(2), 137–154. 10.1016/S0301-0511(00)00076-4

56. Jaušovec, N., & Jaušovec, K. (2005). Differences in induced gamma and upper alpha oscillations in the human brain related to verbal/performance and emotional intelligence. International Journal of Psychophysiology, 56(3), 223–235. 10.1016/j.ijpsycho.2004.12.005

57. Jha, A. P., Morrison, A. B., Dainer-Best, J., Parker, S., Rostrup, N., & Stanley, E. A. (2015). Minds “At Attention”: Mindfulness Training Curbs Attentional Lapses in Military Cohorts. PLOS ONE, 10(2), e0116889. 10.1371/journal.pone.0116889

58. Jo, H.-G., Wittmann, M., Hinterberger, T., & Schmidt, S. (2014). The readiness potential reflects intentional binding. Frontiers in Human Neuroscience, 8. 10.3389/fnhum.2014.00421

59. Karakaş, S. (2020). A review of theta oscillation and its functional correlates. International Journal of Psychophysiology, 157, 82–99. 10.1016/j.ijpsycho.2020.04.008

60. Kasamatsu, A., & Hirai, T. (1966). An electroencephalographic study on the zen meditation (Zazen). Folia Psychiatrica Et Neurologica Japonica, 20(4), 315–336. 10.1111/j.1440-1819.1966.tb02646.x

61. Kassambra, A. (2023). *ggpubr: “ggplot2” Based Publication Ready Plots.* (R package version 0.6.0) [Computer software]. https://rpkgs.datanovia.com/ggpubr/

62. Kayser, C., & Ermentrout, B. (2010). Complex Times for Earthquakes, Stocks, and the Brain’s Activity. Neuron, 66(3), 329–331. 10.1016/j.neuron.2010.04.039

63. Kelly, S. P., Gomez-Ramirez, M., & Foxe, J. J. (2009). The strength of anticipatory spatial biasing predicts target discrimination at attended locations: A high-density EEG study. European Journal of Neuroscience, 30(11), 2224–2234. 10.1111/j.1460-9568.2009.06980.x

64. Keng, S.-L., Smoski, M. J., & Robins, C. J. (2011). Effects of Mindfulness on Psychological Health: A Review of Empirical Studies. Clinical Psychology Review, 31(6), 1041–1056. 10.1016/j.cpr.2011.04.006

65. Khoury, B., Lecomte, T., Fortin, G., Masse, M., Therien, P., Bouchard, V., Chapleau, M.-A., Paquin, K., & Hofmann, S. G. (2013). Mindfulness-based therapy: A comprehensive meta-analysis. Clinical Psychology Review, 33(6), 763–771. 10.1016/j.cpr.2013.05.005

66. Kiken, L. G., Garland, E. L., Bluth, K., Palsson, O. S., & Gaylord, S. A. (2015). From a state to a trait: Trajectories of state mindfulness in meditation during intervention predict changes in trait mindfulness. Personality and Individual Differences, 81, 41–46. 10.1016/j.paid.2014.12.044

67. Klimesch, W. (1999). EEG alpha and theta oscillations reflect cognitive and memory performance: A review and analysis. Brain Research Reviews, 29(2), 169–195. 10.1016/S0165-0173(98)00056-3

68. Klimesch, W. (2012). Alpha-band oscillations, attention, and controlled access to stored information. Trends in Cognitive Sciences, 16(12), 606–617. 10.1016/j.tics.2012.10.007

69. Klimesch, W., Doppelmayr, M., Pachinger, T., & Ripper, B. (1997). Brain oscillations and human memory: EEG correlates in the upper alpha and theta band. Neuroscience Letters, 238(1–2), 9–12. 10.1016/S0304-3940(97)00771-4

70. Klimesch, W., Sauseng, P., & Hanslmayr, S. (2007). EEG alpha oscillations: The inhibition–timing hypothesis—ScienceDirect. 53, 63–88. 10.1016/j.brainresrev.2006.06.003

71. Kobal, G., Wandhöfer, A., & Plattig, K. H. (1975). EEG Power Spectra and Auditory Evoked-Potentials in Transcendental Meditation (TM). 359, R96.

72. Korotkova, T., Ponomarenko, A., Monaghan, C. K., Poulter, S. L., Cacucci, F., Wills, T., Hasselmo, M. E., & Lever, C. (2018). Reconciling the different faces of hippocampal theta: The role of theta oscillations in cognitive, emotional and innate behaviors. Neuroscience and Biobehavioral Reviews, 85, 65–80. 10.1016/j.neubiorev.2017.09.004

73. Kuznetsova, A., Brockhoff, P. B., & Christensen, R. H. B. (2017). lmerTest Package: Tests in Linear Mixed Effects Models *| Journal of Statistical Software*. 82(13), 1–26. 10.18637/jss.v082.i13

74. Lagopoulos, J., Xu, J., Rasmussen, I., Vik, A., Malhi, G. S., Eliassen, C. F., Arntsen, I. E., Saether, J. G., Hollup, S., Holen, A., Davanger, S., & Ellingsen, Ø. (2009). Increased theta and alpha EEG activity during nondirective meditation. *Journal of Alternative and Complementary Medicine (New York*, N.Y.), 15(11), 1187–1192. 10.1089/acm.2009.0113

75. Lee, D. J., Kulubya, E., Goldin, P., Goodarzi, A., & Girgis, F. (2018). Review of the Neural Oscillations Underlying Meditation. Frontiers in Neuroscience, 12. 10.3389/fnins.2018.00178

76. Leicht, G., Troschütz, S., Andreou, C., Karamatskos, E., Ertl, M., Naber, D., & Mulert, C. (2013). Relationship between Oscillatory Neuronal Activity during Reward Processing and Trait Impulsivity and Sensation Seeking. PLOS ONE, 8(12), e83414. 10.1371/journal.pone.0083414

77. Lendner, J. D., Helfrich, R. F., Mander, B. A., Romundstad, L., Lin, J. J., Walker, M. P., Larsson, P. G., & Knight, R. T. (2020). An electrophysiological marker of arousal level in humans. eLife, 9. 10.7554/eLife.55092

78. Libet, B., Gleason, C. A., Wright, E. W., & Pearl, D. K. (1983). Time of conscious intention to act in relation to onset of cerebral activity (readiness-potential). The unconscious initiation of a freely voluntary act. Brain: A Journal of Neurology, 106 (*Pt 3*), 623–642. 10.1093/brain/106.3.623

79. Liebherr, M., Corcoran, A. W., Alday, P. M., Coussens, S., Bellan, V., Howlett, C. A., Immink, M. A., Kohler, M., Schlesewsky, M., & Bornkessel-Schlesewsky, I. (2021). EEG and behavioral correlates of attentional processing while walking and navigating naturalistic environments. Scientific Reports, 11(1), 22325. 10.1038/s41598-021-01772-8

80. Lodha, S., & Gupta, R. (2022). Mindfulness, Attentional Networks, and Executive Functioning: A Review of Interventions and Long-Term Meditation Practice. Journal of Cognitive Enhancement, 6(4), 531–548. 10.1007/s41465-022-00254-7

81. Lomas, T., Edginton, T., Cartwright, T., & Ridge, D. (2014). Men developing emotional intelligence through meditation? Integrating narrative, cognitive and electroencephalography (EEG) evidence. Psychology of Men & Masculinity, 15(2), 213–224. 10.1037/a0032191

82. Lomas, T., Ivtzan, I., & Fu, C. H. Y. (2015). A systematic review of the neurophysiology of mindfulness on EEG oscillations. Neuroscience & Biobehavioral Reviews, 57, 401–410. 10.1016/j.neubiorev.2015.09.018

83. Luck, S. J. (2014). *An Introduction to the Event-Related Potential Technique*. MIT Press. http://ebookcentral.proquest.com/lib/unisa/detail.action?docID=3339822

84. Lüdecke, D. (2018). ggeffects: Tidy Data Frames of Marginal Effects from Regression Models. Journal of Open Source Software, 3(26), 772. 10.21105/joss.00772

85. Lüdecke, D., Ben-Shachar, M. S., Patil, I., Waggoner, P., & Makowski, D. (2021). performance: An R Package for Assessment, Comparison and Testing of Statistical Models. Journal of Open Source Software, 6(60), 3139. 10.21105/joss.03139

86. Lui, K. K., Nunez, M. D., Cassidy, J. M., Vandekerckhove, J., Cramer, S. C., & Srinivasan, R. (2021). Timing of readiness potentials reflect a decision-making process in the human brain. Computational brain & behavior, 4, 264–283. 10.1007%2Fs42113-020-00097-5

87. Lutz, A., Greischar, L. L., Rawlings, N. B., Ricard, M., & Davidson, R. J. (2004). Long-term meditators self-induce high-amplitude gamma synchrony during mental practice. Proceedings of the National Academy of Sciences of the United States of America, 101(46), 16369–16373. 10.1073/pnas.0407401101

88. Lutz, A., Slagter, H. A., Dunne, J. D., & Davidson, R. J. (2008). Attention regulation and monitoring in meditation. Trends in Cognitive Sciences, 12(4), 163–169. 10.1016/j.tics.2008.01.005

89. MacGregor-Fors, I., & Payton, M. (2013). Contrasting Diversity Values: Statistical Inferences Based on Overlapping Confidence Intervals. PloS One, 8, e56794. 10.1371/journal.pone.0056794

90. Magosso, E., De Crescenzio, F., Ricci, G., Piastra, S., & Ursino, M. (2019). EEG Alpha Power Is Modulated by Attentional Changes during Cognitive Tasks and Virtual Reality Immersion. Computational Intelligence and Neuroscience, 2019, 7051079. 10.1155/2019/7051079

91. Malinowski, P. (2013). Neural mechanisms of attentional control in mindfulness meditation. Frontiers in Neuroscience, 7. 10.3389/fnins.2013.00008

92. Manning, J. R., Jacobs, J., Fried, I., & Kahana, M. J. (2009). Broadband Shifts in Local Field Potential Power Spectra Are Correlated with Single-Neuron Spiking in Humans. Journal of Neuroscience, 29(43), 13613–13620. 10.1523/JNEUROSCI.2041-09.2009

93. Matusz, P. J., Dikker, S., Huth, A. G., & Perrodin, C. (2019). Are We Ready for Real-world Neuroscience? Journal of Cognitive Neuroscience, 31(3), 327–338. 10.1162/jocn_e_01276

94. McEvoy, T. M., Frumkin, L. R., & Harkins, S. W. (1980). Effects of meditation on brainstem auditory evoked potentials. The International Journal of Neuroscience, 10(2–3), 165–170. 10.3109/00207458009160494

95. Michailovs, S., Pond, S., Schmitt, M., Irons, J., Stoker, M., Visser, T. A. W., Huf, S., & Loft, S. (2021). The Impact of Information Integration in a Simulation of Future Submarine Command and Control: Human Factors. 10.1177/00187208211045872

96. Miller, K. J., Sorensen, L. B., Ojemann, J. G., & Nijs, M. den. (2009). Power-Law Scaling in the Brain Surface Electric Potential. PLOS Computational Biology, 5(12), e1000609. 10.1371/journal.pcbi.1000609

97. Milz, P., Faber, P. L., Lehmann, D., Kochi, K., & Pascual-Marqui, R. D. (2014). sLORETA intracortical lagged coherence during breath counting in meditation-naïve participants. Frontiers in Human Neuroscience, 8. https://www.frontiersin.org/articles/10.3389/fnhum.2014.00303

98. Mölle, M., Marshall, L., Lutzenberger, W., Pietrowsky, R., Fehm, H. L., & Born, J. (1996). Enhanced dynamic complexity in the human EEG during creative thinking. Neuroscience Letters, 208(1), 61–64. 10.1016/0304-3940(96)12539-8

99. Moore, A. W., Gruber, T., Derose, J., & Malinowski, P. (2012). Regular, brief mindfulness meditation practice improves electrophysiological markers of attentional control. Frontiers in Human Neuroscience, 6. 10.3389/fnhum.2012.00018

100. Morse, D. R., Martin, J. S., Furst, M. L., & Dubin, L. L. (1977). A physiological and subjective evaluation of meditation, hypnosis, and relaxation. Psychosomatic Medicine, 39(5), 304–324. 10.1097/00006842-197709000-00004

101. Neo, P. S.-H., & McNaughton, N. (2011). Frontal theta power linked to neuroticism and avoidance. *Cognitive, Affective*, & Behavioral Neuroscience, 11(3), 396–403. 10.3758/s13415-011-0038-x

102. Ostlund, B. D., Alperin, B. R., Drew, T., & Karalunas, S. L. (2021). Behavioral and cognitive correlates of the aperiodic (1/f-like) exponent of the EEG power spectrum in adolescents with and without ADHD. Developmental Cognitive Neuroscience, 48, 100931. 10.1016/j.dcn.2021.100931

103. Ouyang, G., Hildebrandt, A., Schmitz, F., & Herrmann, C. S. (2020). Decomposing alpha and 1/f brain activities reveals their differential associations with cognitive processing speed. NeuroImage, 205, 116304. 10.1016/j.neuroimage.2019.116304

104. Pathania, A., Schreiber, M., Miller, M. W., Euler, M. J., & Lohse, K. R. (2021). Exploring the reliability and sensitivity of the EEG power spectrum as a biomarker. International Journal of Psychophysiology, 160, 18–27. 10.1016/j.ijpsycho.2020.12.002

105. Paty, J., Benezech, M., Eschapasse, P., & Noël, B. (1978). Neurophysiological study of 47, XYY and 47, XXY psychopaths: Contingent negative variation, evoked potentials and motor nerve conduction. Neuropsychobiology, 4(6), 321–327. 10.1159/000117647

106. Pertermann, M., Mückschel, M., Adelhöfer, N., Ziemssen, T., & Beste, C. (2019). On the interrelation of 1/f neural noise and norepinephrine system activity during motor response inhibition. Journal of Neurophysiology, 121(5), 1633–1643. 10.1152/jn.00701.2018

107. Peterson, E. J., Rosen, B. Q., Campbell, A. M., Belger, A., & Voytek, B. (2017). 1/ f neural noise is a better predictor of schizophrenia than neural oscillations [Preprint]. Neuroscience. 10.1101/113449

108. Pfurtscheller, G., & Lopes da Silva, F. H. (1999). Event-related EEG/MEG synchronization and desynchronization: Basic principles. Clinical Neurophysiology, 110(11), 1842–1857. 10.1016/S1388-2457(99)00141-8

109. Puma, S., Matton, N., Paubel, P.-V., Raufaste, É., & El-Yagoubi, R. (2018). Using theta and alpha band power to assess cognitive workload in multitasking environments. International Journal of Psychophysiology, 123, 111–120. 10.1016/j.ijpsycho.2017.10.004

110. Putman, P., van Peer, J., Maimari, I., & van der Werff, S. (2010). EEG theta/beta ratio in relation to fear-modulated response-inhibition, attentional control, and affective traits. Biological Psychology, 83(2), 73–78. 10.1016/j.biopsycho.2009.10.008

111. Putman, P., Verkuil, B., Arias-Garcia, E., Pantazi, I., & van Schie, C. (2014). EEG theta/beta ratio as a potential biomarker for attentional control and resilience against deleterious effects of stress on attention. Cognitive, Affective, & Behavioral Neuroscience, 14(2), 782–791. 10.3758/s13415-013-0238-7

112. R Core Team. (2020). R: A language and environment for statistical computing. R Foundation for Statistical Computing. https://www.R-project.org/

113. Raufi, B., & Longo, L. (2022). An Evaluation of the EEG Alpha-to-Theta and Theta-to-Alpha Band Ratios as Indexes of Mental Workload. Frontiers in Neuroinformatics, 16. 10.3389/fninf.2022.861967

114. Renard, Y., Lotte, F., Gibert, G., Congedo, M., Maby, E., Delannoy, V., Bertrand, O., & Lécuyer, A. (2010). OpenViBE: An Open-Source Software Platform to Design, Test, and Use Brain– Computer Interfaces in Real and Virtual Environments. Presence, 19(1), 35–53. Presence. 10.1162/pres.19.1.35

115. Rodriguez-Larios, J., Bracho Montes de Oca, E. A., & Alaerts, K. (2021). The EEG spectral properties of meditation and mind wandering differ between experienced meditators and novices. NeuroImage, 245, 118669. 10.1016/j.neuroimage.2021.118669

116. Rodriguez-Larios, J., Faber, P., Achermann, P., Tei, S., & Alaerts, K. (2020). From thoughtless awareness to effortful cognition: Alpha - theta cross-frequency dynamics in experienced meditators during meditation, rest and arithmetic. Scientific Reports, 10(1), 5419. 10.1038/s41598-020-62392-2

117. Rohrbaugh, J. W., & Gaillard, A. W. K. (1983). 13 Sensory and Motor Aspects of the Contingent Negative Variation. In A. W. K. Gaillard & W. Ritter (Eds.), Advances in Psychology (Vol. 10, pp. 269–310). North-Holland. 10.1016/S0166-4115(08)62044-0

118. Rohrbaugh, J. W., Syndulko, K., & Lindsley, D. B. (1976). Brain wave components of the contingent negative variation in humans. *Science (New York*, N.Y.), 191(4231), 1055–1057. 10.1126/science.1251217

119. Schad, D., Vasishth, S., Hohenstein, S., & Kliegl, R. (2020). How to capitalize on a priori contrasts in linear (mixed) models: A tutorial. Journal of Memory and Language, 110, 104038. 10.1016/j.jml.2019.104038

120. Schaworonkow, N., & Voytek, B. (2021). Longitudinal changes in aperiodic and periodic activity in electrophysiological recordings in the first seven months of life. Developmental Cognitive Neuroscience, 47, 100895. 10.1016/j.dcn.2020.100895

121. Schlegel, A., Alexander, P., Sinnott-Armstrong, W., Roskies, A., Tse, P. U., & Wheatley, T. (2013). Barking up the wrong free: Readiness potentials reflect processes independent of conscious will. Experimental Brain Research, 229(3), 329–335. 10.1007/s00221-013-3479-3

122. Schurger, A., Hu, P. “Ben,” Pak, J., & Roskies, A. L. (2021). What Is the Readiness Potential? Trends in Cognitive Sciences, 25(7), 558–570. 10.1016/j.tics.2021.04.001

123. Seager, M. A., Johnson, L. D., Chabot, E. S., Asaka, Y., & Berry, S. D. (2002). Oscillatory brain states and learning: Impact of hippocampal theta-contingent training. Proceedings of the National Academy of Sciences of the United States of America, 99(3), 1616–1620. 10.1073/pnas.032662099

124. Sheehan, T. C., Sreekumar, V., Inati, S. K., & Zaghloul, K. A. (2018). Signal Complexity of Human Intracranial EEG Tracks Successful Associative-Memory Formation across Individuals. Journal of Neuroscience, 38(7), 1744–1755. 10.1523/JNEUROSCI.2389-17.2017

125. Sonkusare, S., Breakspear, M., & Guo, C. (2019). Naturalistic Stimuli in Neuroscience: Critically Acclaimed. Trends in Cognitive Sciences, 23(8), 699–714. 10.1016/j.tics.2019.05.004

126. Takahashi, T., Murata, T., Hamada, T., Omori, M., Kosaka, H., Kikuchi, M., Yoshida, H., & Wada, Y. (2005). Changes in EEG and autonomic nervous activity during meditation and their association with personality traits. International Journal of Psychophysiology: Official Journal of the International Organization of Psychophysiology, 55(2), 199–207. 10.1016/j.ijpsycho.2004.07.004

127. Tanaka, G. K., Peressutti, C., Teixeira, S., Cagy, M., Piedade, R., Nardi, A. E., Ribeiro, P., & Velasques, B. (2014). Lower trait frontal theta activity in mindfulness meditators. Arquivos de Neuro-Psiquiatria, 72, 687–693. 10.1590/0004-282X20140133

128. Tang, Y.-Y., Hölzel, B. K., & Posner, M. I. (2015). The neuroscience of mindfulness meditation. Nature Reviews Neuroscience, 16(4), 213–225. 10.1038/nrn3916

129. Telles, S., Joseph, C., Venkatesh, S., & Desiraju, T. (1993). Alterations of auditory middle latency evoked potentials during yogic consciously regulated breathing and attentive state of mind. International Journal of Psychophysiology, 14(3), 189–198. 10.1016/0167-8760(93)90033-L

130. Telles, S., Nagarathna, R., Nagendra, H. R., & Desiraju, T. (1994). Alterations in auditory middle latency evoked potentials during meditation on a meaningful symbol—“Om”. International Journal of Neuroscience, 76(1–2), 87–93. 10.3109/00207459408985995

131. Telles, S., & Naveen, K. V. (2004). Changes in middle latency auditory evoked potentials during meditation. Psychological Reports, 94(2), 398–400. 10.2466/pr0.94.2.398-400

132. Telles, S., Singh, D., Naveen, K. V., Pailoor, S., Singh, N., & Pathak, S. (2019). P300 and Heart Rate Variability Recorded Simultaneously in Meditation. Clinical EEG and Neuroscience, 50(3), 161–171. 10.1177/1550059418790717

133. Teper, R., & Inzlicht, M. (2013). Meditation, mindfulness and executive control: The importance of emotional acceptance and brain-based performance monitoring. Social Cognitive and Affective Neuroscience, 8(1), 85–92. 10.1093/scan/nss045

134. Teplan, M., Krakovská, A., & Špajdel, M. (2014). Spectral EEG Features of a Short Psycho-physiological Relaxation. Measurement Science Review, 14(4), 237–242. 10.2478/msr-2014-0032

135. Thut, G., Nietzel, A., Brandt, S. A., & Pascual-Leone, A. (2006). Alpha-band electroencephalographic activity over occipital cortex indexes visuospatial attention bias and predicts visual target detection. The Journal of Neuroscience: The Official Journal of the Society for Neuroscience, 26(37), 9494–9502. 10.1523/JNEUROSCI.0875-06.2006

136. Tran, T. T., Rolle, C. E., Gazzaley, A., & Voytek, B. (2020). Linked Sources of Neural Noise Contribute to Age-related Cognitive Decline. Journal of Cognitive Neuroscience, 32(9), 1813–1822. 10.1162/jocn_a_01584

137. Travers, E., & Haggard, P. (2021). The Readiness Potential reflects the internal source of action, rather than decision uncertainty. European Journal of Neuroscience, 53(5), 1533–1544. 10.1111/ejn.15063

138. Travis, F., Tecce, J., Arenander, A., & Wallace, R. K. (2002). Patterns of EEG coherence, power, and contingent negative variation characterize the integration of transcendental and waking states. Biological Psychology, 61(3), 293–319. 10.1016/s0301-0511(02)00048-0

139. Tsai, J.-F., Jou, S.-H., Cho, W., & Lin, C.-M. (2013). Electroencephalography when meditation advances: A case-based time-series analysis. Cognitive Processing, 14(4), 371–376. 10.1007/s10339-013-0563-3

140. Tye, C., Rijsdijk, F., & McLoughlin, G. (2014). Genetic overlap between ADHD symptoms and EEG theta power. Brain and Cognition, 87, 168–172. 10.1016/j.bandc.2014.03.010

141. Vallat, R., & Walker, M. P. (2021). An open-source, high-performance tool for automated sleep staging. eLife, 10, e70092. 10.7554/eLife.70092

142. Virk, T., Letendre, T., & Pathman, T. (2024). The convergence of naturalistic paradigms and cognitive neuroscience methods to investigate memory and its development. Neuropsychologia, 196, 108779. 10.1016/j.neuropsychologia.2023.108779

143. von Stein, A., & Sarnthein, J. (2000). Different frequencies for different scales of cortical integration: From local gamma to long range alpha/theta synchronization. International Journal of Psychophysiology: Official Journal of the International Organization of Psychophysiology, 38(3), 301–313. 10.1016/s0167-8760(00)00172-0

144. Wang, P., He, Y., Maess, B., Yue, J., Chen, L., Brauer, J., Friederici, A. D., & Knösche, T. R. (2022). Alpha power during task performance predicts individual language comprehension. NeuroImage, 260, 119449. 10.1016/j.neuroimage.2022.119449

145. Wang, P., Knösche, T. R., Chen, L., Brauer, J., Friederici, A. D., & Maess, B. (2021). Functional brain plasticity during L1 training on complex sentences: Changes in gamma-band oscillatory activity. Human Brain Mapping, 42(12), 3858–3870. 10.1002/hbm.25470

146. Wickham, H. (2016). ggplot2: Elegant Graphics for Data Analysis. Springer-Verlag New York. https://ggplot2.tidyverse.org.

147. Wickham, H., Averick, M., Bryan, J., Chang, W., McGowan, L., François, R., Grolemund, G., Hayes, A., Henry, L., Hester, J., Kuhn, M., Pedersen, T., Miller, E., Bache, S., Müller, K., Ooms, J., Robinson, D., Seidel, D., Spinu, V., … Yutani, H. (2019). Welcome to the Tidyverse. Journal of Open Source Software, 4(43), 1686. 10.21105/joss.01686

148. Wong, K. F., Teng, J., Chee, M. W. L., Doshi, K., & Lim, J. (2018). Positive effects of mindfulness-based training on energy maintenance and the EEG correlates of sustained attention in a cohort of nurses. Frontiers in Human Neuroscience, 12. 10.3389/fnhum.2018.00080

149. Yoshida, K., Takeda, K., Kasai, T., Makinae, S., Murakami, Y., Hasegawa, A., & Sakai, S. (2020). Focused attention meditation training modifies neural activity and attention: Longitudinal EEG data in non-meditators. Social Cognitive and Affective Neuroscience, 15(2), 215–224. 10.1093/scan/nsaa020

150. Zhang, W., Zheng, R., Zhang, B., Yu, W., & Shen, X. (1993). An Observation on Flash Evoked Cortical Potentials and Qigong Meditation. The American Journal of Chinese Medicine, 21(03n04), 243–249. 10.1142/S0192415X93000285

151. Zietsch, B. P., Hansen, J. L., Hansell, N. K., Geffen, G. M., Martin, N. G., & Wright, M. J. (2007). Common and specific genetic influences on EEG power bands delta, theta, alpha, and beta. Biological Psychology, 75(2), 154–164. 10.1016/j.biopsycho.2007.01.004

